# The mechanism of biofilm degradation by a detachable tailspike of gene transfer agents

**DOI:** 10.64898/2026.07.19.739414

**Authors:** Pavol Bardy, Phuong M. Nguyen, Yan Liu, Matthew W. Craske, Nicholas Read, Rebecca M. Davies, Johan P. Turkenburg, Samuel J. Hart, Alfred A. Antson, Paul C. M. Fogg

**Affiliations:** York Structural Biology Laboratory, Department of Chemistry, University of York, Heslington, United Kingdom; York Biomedical Research Institute, University of York, York, United Kingdom; Division of Cryo-EM and Bioimaging, SSRL, SLAC National Accelerator Laboratory, Stanford University, Menlo Park, California, USA; Department of Biology, University of York, York, United Kingdom

## Abstract

Gene transfer agents (GTAs) are phage-derived elements that have evolved repeatedly across diverse prokaryotes, where they drive high-frequency horizontal gene transfer (HGT). Here, we demonstrate that the *Rhodobacter capsulatus* GTA (RcGTA) tailspike protein, TspA, is a potent biofilm-degrading enzyme. Purified TspA is effective at both preventing initial biofilm formation and clearing established, mature biofilms. Crucially, TspA enhances RcGTA-mediated gene transfer, suggesting that this enzyme facilitates GTA navigation through the extracellular matrix. Unlike the permanently anchored tailspikes of canonical phages, TspA possesses a unique β-sandwich N-terminal domain that enables its dissociation from mature particles and engages in biofilm polysaccharide recognition. Our findings indicate that TspA is an evolutionary adaptation used by GTAs to optimize HGT within complex, densely packed microbial biofilm communities.

## INTRODUCTION

Bacteria show a remarkable ability to harness junk or even harmful genetic material for their own benefit. Examples of this are gene transfer agents (GTAs), “tamed” prophages that transfer random segments of bacterial DNA between closely related cells (1). GTAs have evolved independently in several prokaryotic species, are vertically inherited, and typically persist for millions of years. GTAs are released by lytic burst yet, despite this cost to the individual producer, there must be a strong fitness benefit to the rest of the population. It has been shown that some GTA genes are beneficial under metabolic stress (2, 3) and at the molecular level, GTA-mediated gene transfer increases population fitness by repairing deleterious mutations in recipients (4). As a consequence of this transfer, virulence factors and antimicrobial genes are transmitted by GTAs at exceptionally high rates in natural environments (5, 6).

The fitness cost of GTA production is relatively high, with up to 3% of the population in *Rhodobacter* (7) and 9% in Bartonella (8) lysing as a result of GTA production and release. Recipients that lose the GTA-producing phenotype should quickly arise and dominate in the environment (9–11). In addition, in a non-constrained aquatic environment, the released particles would likely be rapidly diluted from the producer cultures, further limiting the advantage of their production (10, 11). This highlights a gap in our understanding of how GTA production increases fitness over evolutionary timeframes and is thus maintained in bacterial populations.

Microbial cells often survive in an adhered state covered by a protective biofilm matrix. This matrix is continuously remodulated to allow biofilm maturation and dispersion, ultimately spreading the culture to new niches (12). Recently, several molecular links between *Rhodobacterales* GTA production and biofilm formation have been identified. A serine acetyltransferase CysE1 is required for both optimal RcGTA recipient ability and biofilm production in *Rhodobacter* (13). The second messenger bis-(3′-5′)-cyclic dimeric GMP, which is a conserved regulator of biofilm formation (14), also affects the production of alphaproteobacterial GTAs (15–17). In *Phaeobacter*, a strain deficient in the production of the antibiotic secondary metabolite tropodithietic acid shows increased GTA production and forms biofilm faster than the wild type (18). Moreover, 15 gene families predicted to be involved in biofilm formation were shown to co-evolve with *Rhodobacterales* GTA genes (9), suggesting that the fitness advantage of GTAs may arise in a biofilm lifestyle.

The virion of *Rhodobacterales* GTA resembles that of a long flexible-tailed phage with the Siphovirus morphology. However, it differs in possessing an oblate head and an unusually short tail (19). Proteins forming the virion are encoded in several genomic loci (20, 21), which prevents the GTA from packaging its entire structural locus and thus reduces the possibility of reverting to a phage lifestyle. This fragmented organisation also creates an opportunity for some GTA genes to evolve more rapidly compared to the conserved virion gene cluster (22, 23). In *Rhodobacter capsulatus*, the ectopic GTA loci encode a variety of receptor-binding proteins, namely head spikes and tail fibres, as well as the lytic module. The head spikes facilitate the attachment of *R. capsulatus* (Rc) GTAs to the recipient cell capsule, thereby mediating initial host recognition (19, 24). The tail fibres are located at the distal tip of the baseplate and are essential for RcGTA-mediated gene delivery (20). Three other domains protrude from the baseplate core. These are the megatron peripheral TIM-barrel domain, the hub oligosaccharide-binding domain and the distal tail protein insertion domain, all encoded within the conserved virion gene cluster. The exact function of these domains has remained unknown.

In this study, by combining molecular characterization with cryo-electron tomography of intact cultures, we discovered an RcGTA virion-associated component, Rcc02623 (here renamed as TspA), as a potent inhibitor of biofilm formation and mature biofilm-degrading enzyme. TspA forms hexamers of trimers and is loosely associated with the protruding domains of the baseplate as well as the tail tube of the RcGTA particle. The protein is conserved among *Rhodobacterales* GTAs, but no homolog is present among phages harbouring RcGTA-like baseplate. These findings showcase that the *Rhodobacterales* GTAs have evolved differently from related phages to navigate through biofilms and to target cells growing within biofilm-associated communities.

## RESULTS

### TspA is a conserved protein co-produced with RcGTAs

Previous RNAseq data showed that TspA is upregulated in the RcGTA overproducer strain DE442, yet not essential for RcGTA-mediated gene transfer (20, 25). Here, the DE442 strain was grown under conditions that promote RcGTA overproduction and the cell-free supernatant was assessed by shotgun mass spectrometry. Our proteomic analysis showed that TspA was one of the most abundant proteins in the sample, with normalized spectral counts comparable to the normally dominant major capsid protein (**Table S1**).

We next decided to test whether the *tspA* gene (*rcc02623*) is associated with RcGTA production in a clean genetic background, since the DE442 strain was generated by random mutagenesis and carries many RcGTA non-associated changes (26). Instead, for these experiments, we used the wildtype *R. capsulatus* SB1003 strain and two isogenic deletion mutants, one lacking the specific RcGTA repressor protein Rcc00280 (GTA overproducer) (27) and one lacking the specific RcGTA activator protein GafA (GTA-null) (25). We performed RT-qPCR quantification of *rcc02622-24* transcripts in wildtype compared to SB1003Δ*280* and SB1003ΔgafA. The result showed ∼1000-fold increase in the transcription of the *tspA* gene, comparable with the increase in RcGTA major capsid protein transcript and consistent with the proteomic data (**Fig. 1A**). In contrast, the transcripts of the neighbouring genes, *rcc02622 and rcc02624*, showed only a negligible increase in the SB1003Δ*280* overproducer (**Fig. 1A**).

**Fig 1.**
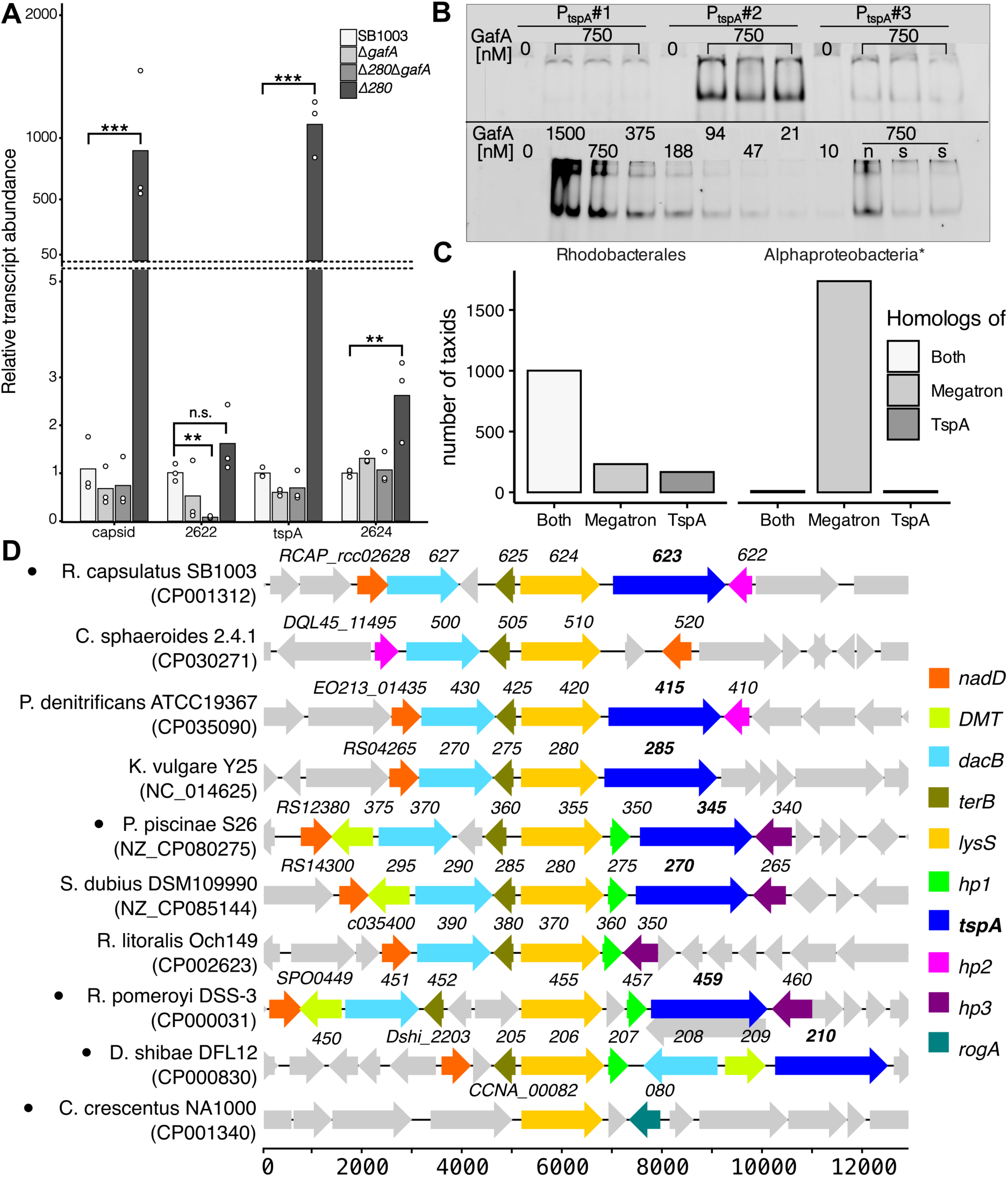
TspA and RcGTA co-regulation and conservation. **A)** Relative expression levels of RcGTA major capsid gene and *rcc02622-02624* within Δ*gafA*, Δ*280*Δ*gafA,* Δ*280* and SB1003 wild-type control. Average relative expression of three biological replicates represented as bars and individual replicates as dots (n=3). Statistical significance of ΔCt values was calculated via one-way ANOVA with Tukey Post-Hoc pairwise multiple comparisons (*** p <0.001, ** p <0.01, * p <0.05 and n.s. p >0.05). **B)** Electromobility shift assay of the *tspA* promoter upon addition of the RcGTA regulator, GafA. Three different regions upstream of *tspA* were tested (top), with the sequence of each region and uncropped gels shown in **Fig. S1**. The observed effect for region PtspA#2 was concentration dependent (bottom). "s", non-fluorescent competitor DNA identical to the target; "n", non-fluorescent competitor DNA that is not expected to bind. **C)** The presence of TspA and megatron homologs in *Rhodobacterales* species and Alphaproteobacteria species, excluding *Rhodobacterales* (*) as identified by PSI-BLAST with hits filtered to coverage >75 %. Raw data containing the hits are present in **Data S1. D)** Conservation of the *tspA* locus in selected representatives of *Rhodobacterales* and distantly related *Caulobacter*.

Apart from the RcGTA cluster, the GafA regulator also binds to its own promoter, positively autoregulating the RcGTA production loop (25). The RT-qPCR analysis showed that upon deletion of GafA, there is a decrease in the production of *tspA* transcripts (**Fig. 1A**). We thus tested if the GafA directly binds to the *tspA* promoter, using an electrophoretic mobility shift assay. We examined individual 50 bp-long segments of the 150bp-long region upstream of the *tspA* start codon (**Fig. S1**). We found that the 50-100 bp upstream region (P_tspA_#2) shifted in the presence of a GafA protein (**Fig. 1B-C**), confirming binding of GafA to this region.

Next, we assessed if TspA is a conserved feature of GTA producers by inspecting how frequently it is encoded in genomes of other alphaproteobacterial GTA-encoding species using PSI-BLAST. We filtered for species which contained both TspA and a GTA marker gene product homolog. As the marker, we selected megatron (Rcc01698), as it is a multidomain protein with no long-query homologs outside of GTA-type baseplates. Almost all *Rhodobacterales* species that encoded megatron also encoded TspA homolog (**Fig. 1C, Data S1**). In contrast, Alphaproteobacteria species, excluding *Rhodobacterales,* encoded the megatron but lacked a TspA homolog (**Fig. 1D, Data S1**). The gene module analysis shows that *tspA* is encoded within the same genome module throughout different *Rhodobacterales* species, although, in some GTA-encoding species such as *Cereibacter sphaeroides* 2.4.1, and *Roseobacter litoralis* Och149, it was lost (**Fig. 1E**). The *lysS* gene (*rcc02624*) was consistently found adjacent to *tspA* and, interestingly, in *Caulobacterales,* which lack *tspA,* it is adjacent to another GTA-associated gene, the regulator *rogA* (**Fig. 1E**) (4). Overall, this shows that *tspA* is a conserved feature of *Rhodobacterales* GTA-producing strains, likely arising in the early stages of the order’s divergence.

### Addition of TspA increases gene transfer rates and promotes biofilm degradation

To assess whether TspA influences RcGTA-mediated gene transfer, we compared the efficiency of RcGTAs produced by an overproducer strain against an isogenic Δ*tspA* deletion strain. When recipient cultures were grown under aerobic conditions, there was no significant difference in the frequency of gene transfer events between the two groups. However, when recipient cultures were grown anaerobically, Δ*tspA* RcGTAs produced 70% fewer gene-transfer events than the *tspA+* control (**Fig. 2A**). This reduction was mitigated when purified TspA was added to the Δ*tspA* RcGTAs prior to the assay. This suggests TspA directly affects the efficiency of RcGTA gene transfer.

**Fig 2.**
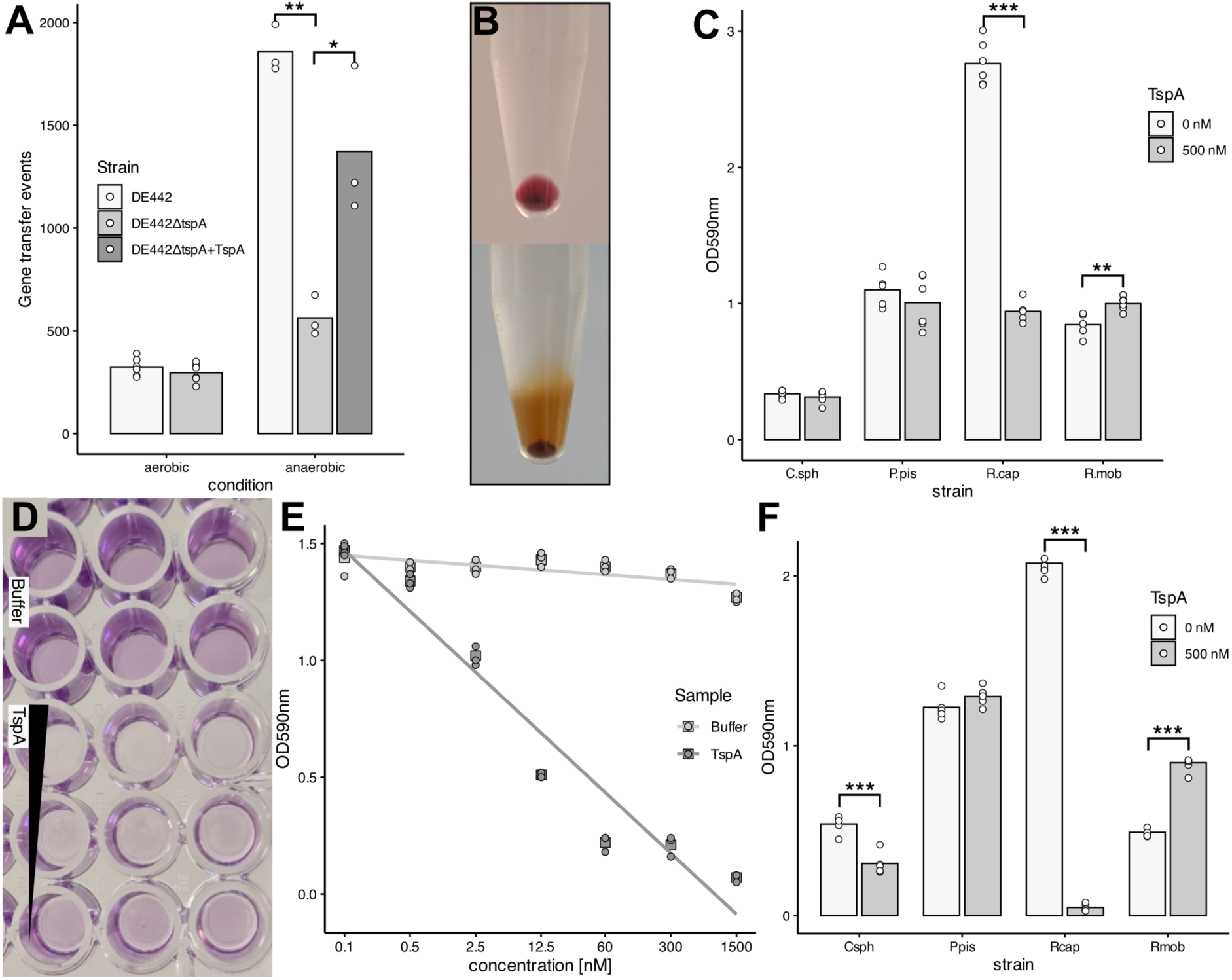
Effect of TspA on RcGTA-mediated gene transfer and biofilm removal. **A)** RcGTA gene transfer assay. Statistical significance was derived using a paired t-test and one-way ANOVA with Tukey post-hoc pairwise multiple comparisons for aerobic and anaerobic conditions, respectively (**, p <0.005; *, p <0.05; and p >0.05, not shown). **B)** Pellet of R. capsulatus B10 cultures grown in minimal media to late stationary phase and centrifuged at 8000 G for 5 min. The aerobic culture is shown on top and the anaerobic on the bottom. Loose substance above the firm cell pellet corresponds to EPS. **C)** Readings of a residual biofilm formed by different *Rhodobacterales* species upon treatment with buffer or TspA, respectively. Statistical significance was calculated via a paired t-test (***, p<0.001; **, p <0.01; and p >0.05, not shown). **D)** Crystal violet-stained biofilm formed on the polystyrene 96-well plate by R*. capsulatus*. The triangle depicts a decreasing concentration of TspA, correlating with an increasing level of residual biofilm. **E)** The level of biofilm formation by *R. capsulatus* SB1003 upon 72-hour incubation after addition of different concentrations of buffer or TspA at time zero. **F)** The level of biofilm formation by different *Rhodobacterales* species upon 72-hour incubation after addition of buffer or TspA at time zero. Csph, *Cereibacter sphaeroides*; Ppis, *Phaeobacter piscinae*; Rcap, *Rhodobacter capsulatus*; Rmob, *Ruegeria mobilis*. Statistical significance was calculated via paired t-test (***, p<0.001; p >0.05, not shown). The raw data of the experiments are shown in **Data S2**.

Under anaerobic conditions, *R. capsulatus* produces more surface-associated slime polysaccharide (28) as well as extracellular polymeric substance (EPS) that forms the basis of a biofilm matrix (13) (**Fig. 2B**). We thus tested if TspA has any effect on biofilm disposal. *R. capsulatus* SB1003 showed potent biofilm formation when grown in uncoated polystyrene 96-well plates, as estimated by crystal violet staining (29), and we used this strain for subsequent biofilm assays, growing the biofilms for 48 hours. After 2-hour incubation with the 500 nM concentration of protein, we observed 66 % reduction in the residual biofilm of SB1003 (**Fig. 2C**). Biofilms of other tested *Rhodobacterales* species were only weakly affected by the addition of the protein. Next, we tested the prophylactic effect of TspA against biofilm formation, with the protein added to the wells along with the culture at time zero and the mixture incubated for 72 hours. Here, almost complete prevention of biofilm formation was achieved at TspA concentrations as low as 60 nM (**Fig. 2D-F**). Similarly, the homolog of TspA from marine bacterium *Tritonibacter* (former *Ruegeria*) *mobilis,* another model producer of GTAs (5), showed a concentration-dependent activity in the prophylaxis of the host biofilm formation, with a statistically significant prophylactic effect shown also against related marine species, *Phaeobacter piscinae* (**Fig. S3)**. We also expressed *Phaeobacter* TspA homolog; however, we obtained low yields and non-reproducible activity of the purified product (**Fig. S3**), with the protein likely being toxic to the producer *E. coli*.

### TspA is a trimeric multidomain tailspike requiring a calcium for its function

To understand the molecular mechanism of TspA, we purified the native protein directly from *R. capsulatus* **(Fig. S2)** and determined its structure by single-particle cryo-EM analysis, achieving an average resolution of 2.0 Å (**Fig. 3A-F, Fig. S4**). The structure showed that TspA forms a homotrimer, with each subunit composed of four domains (**Fig. 3F**): **1)** an N-terminal β-sandwich domain typical for carbohydrate-binding modules (residues 1-180), **2)** a right-handed β-helix domain (residues 181-336 and 412-674), a fold characteristic of carbohydrate active enzymes (30), containing a conserved PF12218-like cap region (residues 181-285) (31), **3)** an insertion domain (residues 337-411), and **4)** C-terminal domain (residues 675-763). The insertion and C-terminal domains each comprise a six β-strands forming a barrel, often associated with binding to diverse biological polymers (32). Structural similarity search for the TspA region 181-763 detected similar folds in several phage tailspikes (**Table S2**). The N-terminal domain of TspA displays significant structural similarity only to one right-handed β-helix protein, the α-1,3-glucanase produced by *Streptomyces thermodiastaticus* that has been shown to degrade extracellular glycans (HHpred, E-value 3.69E-11, residues 37-669) (33) (**Fig. S5**).

**Fig. 3.**
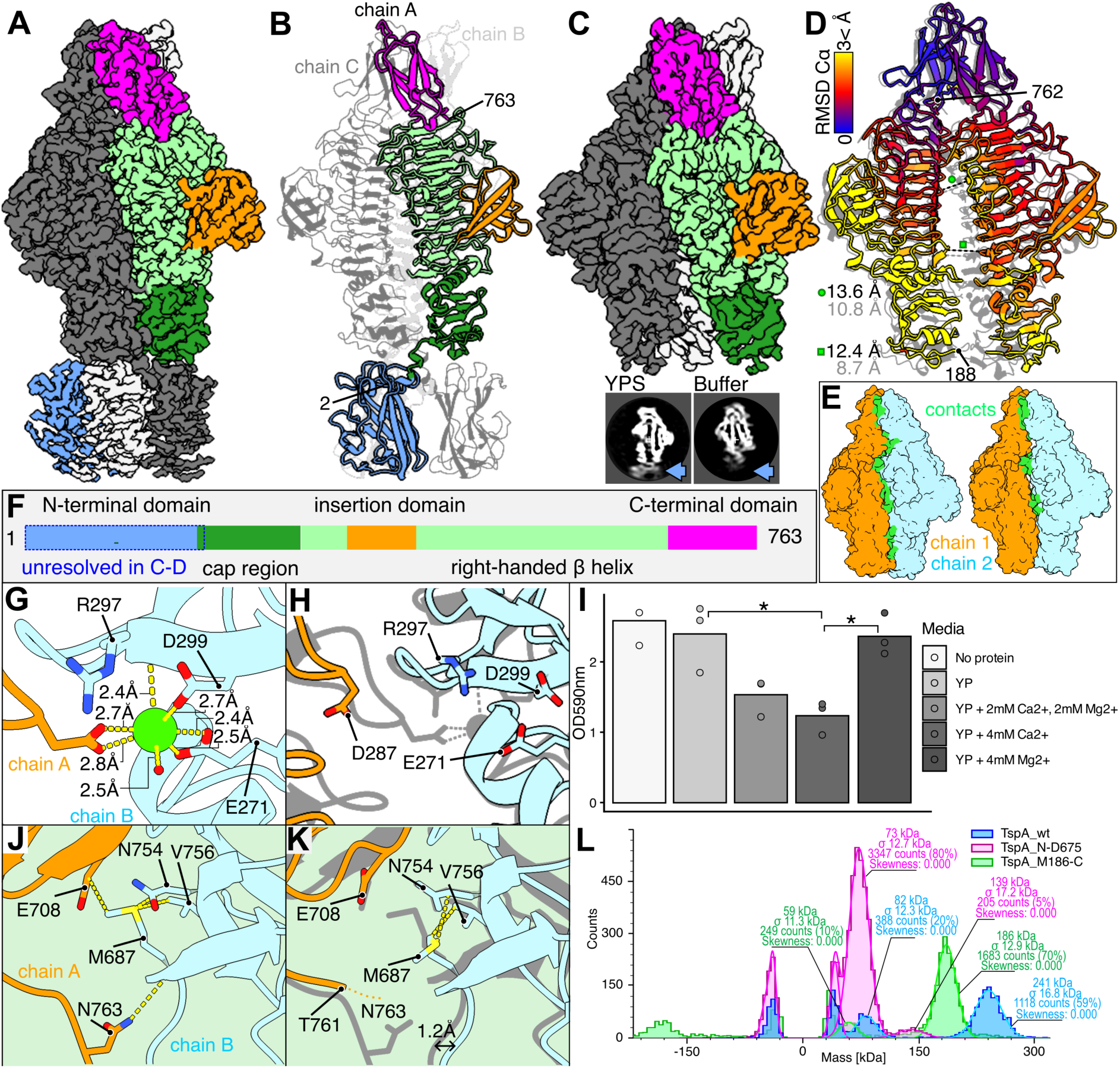
Structure of purified TspA derived from single particle analysis. **A)** Cryo-EM map of the TspA closed state when incubated in the growth media YPS. The map is coloured based on the protein domain depiction, as labelled in panel E. **B)** Protein model built into the map from panel A. **C)** Cryo-EM map of the TspA open state when incubated in buffer. An x-slice projection of maps of TspA incubated in YPS or buffer, respectively. The arrow depicts the position of the N-terminal domain. **D)** Top: superimposition of the models of the closed (grey) and open (coloured based on C_α_ root mean square deviation), aligned based on their C-terminal domains. Distances between two subunits of each state are shown near symbols, black corresponding to the open and grey to the closed conformation. **E)** Interchain contacts of closed (left) and open (right) state of TspA. N-terminal domain is not shown. **F)** Schematic of the protein domain composition. **G-H)** The calcium ion coordination site in the closed (G) and open (H) state (superimposed closed state shown in grey for comparison). Individual residues coordinating the ion (limegreen) are shown in sticks, and distances between the coordinating atoms and the ion are depicted. **I)** The biofilm disposal assay with the reaction incubated in yeast-peptone (YP) media containing different amounts of divalent cations. Bars represent averages of biological triplicate, dots individual measurements. Statistical significance was calculated via one-way ANOVA with Tukey Post-Hoc pairwise multiple comparisons (*, p<0.05). Raw data are present in **Data S2**. **J-K)** The site near M687, showing the major difference in the conformation of the C-terminal domain between the closed (I) and open (J) state (superimposed closed state shown in grey for comparison). Individual residues with distance <4.2 Å from the M687 are shown in sticks, the contacts are shown as lines. **L)** Mass photometry graph, showing mass-related counts for wildtype (blue), N-terminal (M186-C, green) and C-terminal truncations (N-D675, magenta).

We resolved two distinct conformation states of TspA: a closed state with a well-defined N-terminal domain (**Fig. 3A-B**), and an open state in which this domain is unresolved (**Fig. 3C-D**). These states were observed following incubation in rich YPS growth media and in purification buffer (Tris 20 mM, pH=7.5; NaCl 250 mM), respectively. In YPS media, ∼52% of particles lacked a resolved N-terminal domain, exhibiting an architecture similar to the buffer-incubation condition (**Fig. S6**). In the closed state, the cap region is more rigid and the β-helix domains pack closer together (**Fig. 3D, E**), with a calculated solvation free energy of assembly formation, Δ^i^G, of -19.4 kcal/mol per chain, as estimated by PDBePISA (34).

Notably, this conformation coordinates a calcium ion at the inter-subunit interface by Glu271, Asp299 and Arg297 of one subunit and Asp287 of another subunit (**Fig. 3G-H; Fig. S7**), contributing an additional Δ^i^G of -12.3 kcal/mol per chain. In contrast, in the open state with the unresolved N-terminal domains, the inter-subunit contacts are substantially weaker (Δ^i^G of -3.9 kcal/mol) and are mediated primarily by the C-terminal domain (**Fig. 3E**). In the closed state, another calcium ion is coordinated by Asp542, Asp602 and Asn635 of a single subunit (**Fig. S7**), while in the open state, these residues are located to far apart to facilitate coordination. In the absence of calcium, TspA loses its enzymatic activity, which is not restored by the addition of magnesium (**Fig. 3I**). This suggests the calcium-coordinating closed state represents the active state of the protein.

The C-terminal domains of both states share nearly identical structures (RMSD_C_α=0.307), with the only notable difference occurring around Met678. In the open state, Met678 is present in a different conformation, losing hydrophobic contact with Glu708 and a main-chain H-bond with Asn763, both in the neighbouring subunit (**Fig. 3J-K**). This increases the flexibility of Asn763, which loses hydrophobic contacts with Tyr697 and Met672, both from the original subunit, allowing for a more open state of the central β-helical domain. We confirmed the critical role of the C-terminal domain in inter-subunit contacts by deleting it. The truncated variant (1–674) was predominantly monomeric under the tested conditions, whereas the N-terminal truncation did not affect the stoichiometry (**Fig. 3L**).

### The active site of TspA is located at the inter-subunit interface

To shed light on the molecular mechanism of TspA biofilm digestion, we performed targeted mutagenesis of residues in conserved clefts of the central β-helix domain that could serve as active sites. Two such clefts were identified, one located within a single subunit and the other at the interface of two adjacent subunits (**Fig. 4A-C**). Both these sites were shown to bind carbohydrates in distant homologs of TspA (**Fig. S5**). The intradomain cleft contains three conserved glutamic acids (positions Glu377, Glu571 and Glu575) and one aspartic acid (Asp645) (**Fig. 4C**). Individually removing the charge from these residues by performing Glu>Gln and Asp>Asn substitutions did not hamper the activity of the protein in the case of Gln571Q and Asn645N (**Fig. 4F**). Due to instability, in the case of Glu377Q and Glu575Q, a small amount of soluble protein could be obtained only by a crude metal-affinity pull-down (**Fig. S8**). However, even these low-concentration samples reduced biofilm formation in the prophylactic assay (**Fig. 4F**), suggesting that the conserved charged amino acids in the intradomain cleft are not involved in catalysis.

**Fig. 4.**
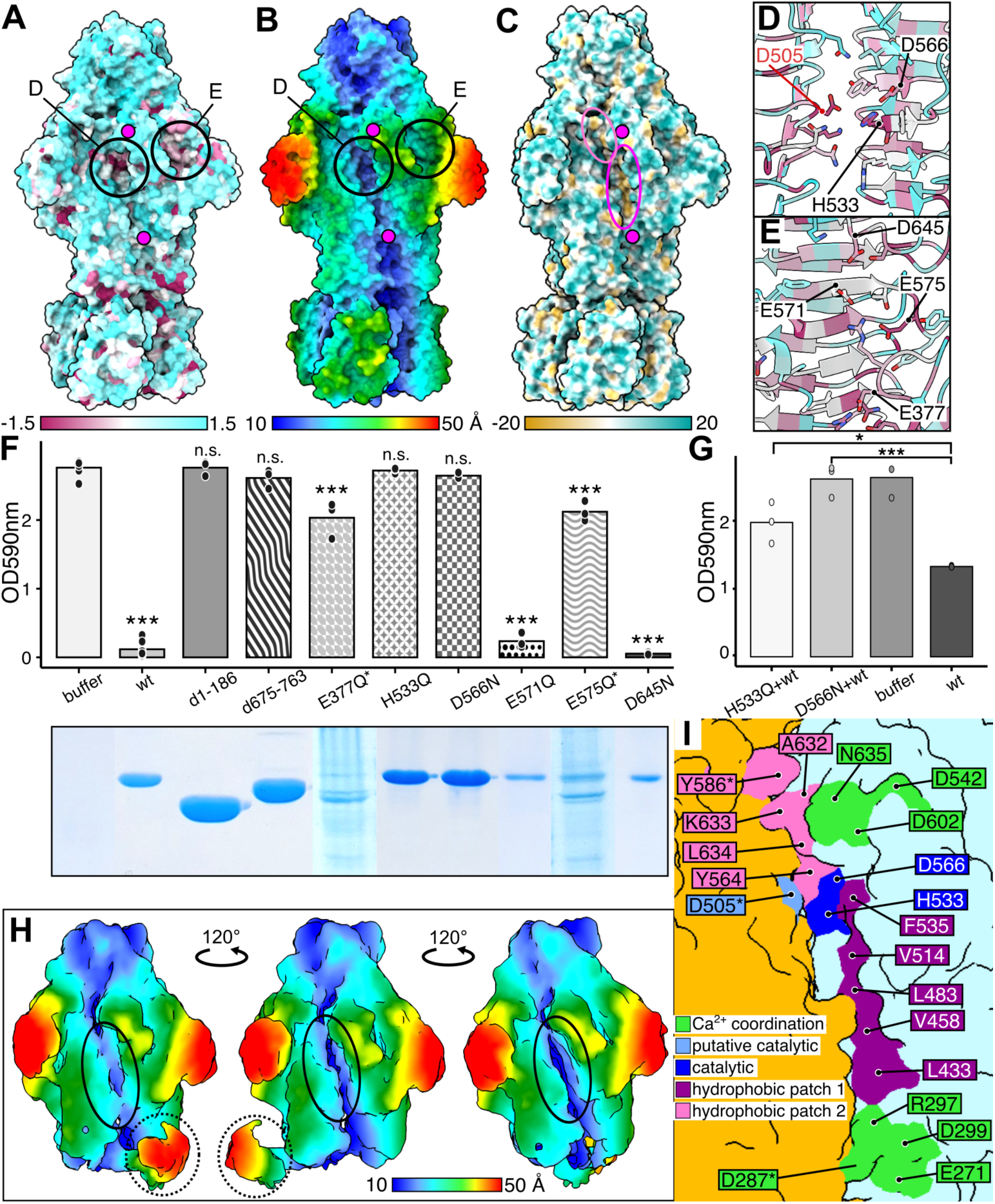
Characterisation of TspA active site. **A-C)** The molecular surface of TspA coloured based on: A) sequence conservation, the values correspond to an entropy-based measure from AL2CO (35), with maroon representing conserved and cyan non-conserved residues; B) cylindrical distance from the central z-axis; C) hydrophobicity, with yellow representing more hydrophobic and cyan more hydrophilic surface. Calcium ions are depicted as magenta circles. The magenta and pink oval highlights hydrophobic patches 1 and 2, respectively. **D-E)** Close-up of the conserved regions of the inter-subunit cleft (D) and intra-subunit cleft (E) with protein shown in ribbon and residues forming the cleft shown as sticks. Putative and experimentally verified catalytic residues are depicted in red and black, respectively. **F)** Top, chart of the biofilm prophylaxis assay with different mutants of TspA added at 500 nM. Bars represent averages, dots individual measurements. Statistical significance was calculated between buffer and individual protein variant (***, p<0.001; n.s, p>0.05). *, unstable protein whose final concentration could not be estimated. Bottom, SDS-PAGE lanes showing individual purified proteins used in the assay. Full gel images are present in **Fig. S8**. **G)** The biofilm disposal assay with 500 nM of wild-type (wt) TspA upon pre-treatment with TspA mutants (***, p<0,001; *, p<0.05). The raw data of the experiments are shown in **Data S2**. **H)** Three views of the asymmetric map of TspA upon incubation with biofilm, showing a bent N-terminal domain (dashed circle) and extra density in the inter-subunit interface (solid circle). The map is coloured according to the cylindrical distance from the central z-axis. **I)** The surface representation of residues putatively involved in ligand binding, coloured based on their function/location. Different chains are coloured orange and light blue, respectively. *, residues from the chain shown in orange.

The interdomain cleft contains conserved Asp505 of the left subunit, and His533 and Asp566 of the right subunit (**Fig. 4B**). Both mutations of Asp566 to Asn, and His533 to Gln, completely ablated the biofilm-disposal activity (**Fig. 4F**), suggesting they are directly involved in the catalytic process. The mutant constructs also protected the biofilm from wild-type TspA activity in a competitive inhibition assay, suggesting that the modified proteins still retain the ability to bind their substrate (**Fig. 4G**). Taken together, we conclude that the active site of the TspA trimer is located in the interdomain interface.

To further elucidate the mechanism of substrate recognition by TspA, we mixed the catalytically dead variant D566N with a resuspended residual biofilm material formed by *R. capsulatus* SB1003 on the surface of the polystyrene plate wells and collected a single-particle dataset. After processing, we observed a major difference in the conformation of the N-terminal domains, with the domains localised around the sides of the cap regions (**Fig. S9**). When reconstructing asymmetrically, we identified a state in which a single N-terminal domain bends and localises near the β-helix domain (**Fig. 4H**), comprising ∼35 % of particles and resembling the topology of the glucanase from *S. thermodiastaticus* (**Fig. S5**).

Interestingly, the inter-subunit interface located clockwise from the bent domain contains extra density (**Fig. 4H**). The density location corresponds to the hydrophobic patch, which lies just below the confirmed active site (hydrophobic patch 1, **Fig. 4C**). Interestingly, there is another hydrophobic patch located just above the active site (hydrophobic patch 2, **Fig. 4C**), but no significant differences in density were observed for this region.

The map anisotropy caused by the preferred orientation in the dataset did not enable us to resolve the extra density further. We also collected a dataset on TspA mixed with the EPS produced by the B10 strain under anaerobic conditions (**Fig. 2B**); however, this data showed an even more pronounced preferred orientation (**Fig. S9**). Nevertheless, as the extra density corresponds to the likely ligand-binding site as identified by mutagenesis, we believe the obtained map represents the active state of the protein with attached biofilm polysaccharide (**Fig. 4I**).

### TspA is loosely attached to RcGTA virions

The previously determined structure of the RcGTA virion did not contain any density corresponding to TspA (19). We therefore imaged RcGTA-producing cultures by cryo-ET to determine whether the protein is associated with RcGTA release. Interestingly, most native virions in this crude sample contained an additional density on their baseplate when compared to purified particles **(Fig. 5A)**. To assess the identity of this density, we performed a subtomogram averaging of the baseplate and found that the extra density matched the shape of TspA trimers, with six trimers arranged around the baseplate core (**Fig. 5B, Fig. S10**).

**Fig. 5.**
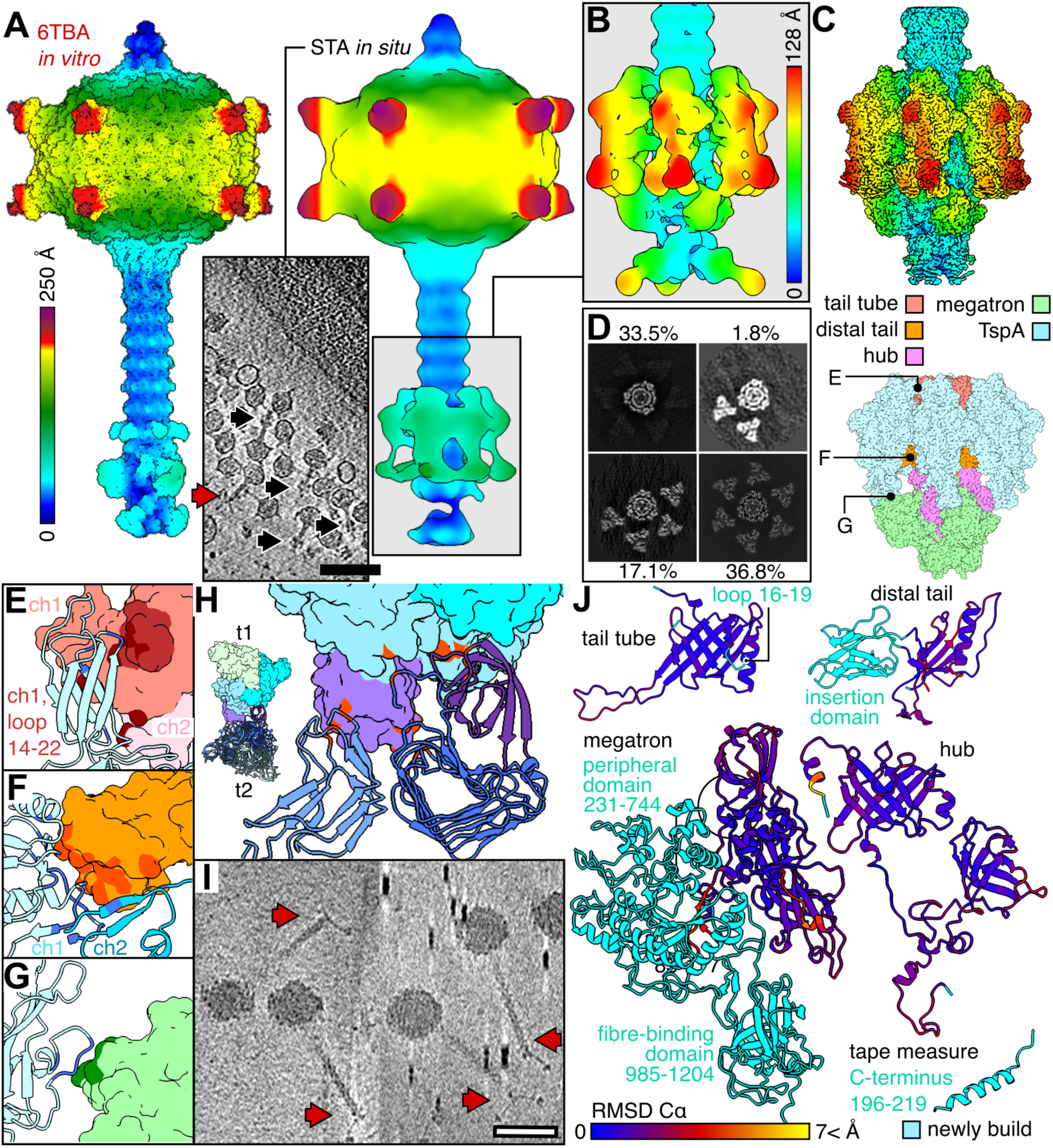
TspA is loosely associated with the RcGTA baseplate. **A)** RcGTA virion surface derived from purified particles (on left) and *in situ* (right). The capsids are coloured according to their cylindrical distance from the central z-axis. An example of the *in situ* tomogram is shown in the middle. Both RcGTAs with a bulky (black arrows) and thin baseplate density (red arrow) were observed. **B)** Localised subtomogram averaging of the baseplate, coloured according to a distance from the central z-axis. **C)** Single particle analysis of the baseplate with attached TspA, the final map is shown coloured based on the cylindrical distance from the central z-axis (top), built protein models are depicted in surface (bottom). **D)** Z-slice projections of baseplate 3D classes, showcasing a different number of attached TspA trimers. The percentage represents the share of all particles; the remaining 10.8 % were junk particles. **E-G)** Interaction interface between TspA and tail tube (E), distal tail (F), and megatron (G). TspA is depicted in ribbons, baseplate proteins in surface. Interacting residues are highlighted in darker shades. If relevant, multiple interacting chains (ch) are shown. **H)** Interaction interface between two trimers (t) of TspA within the baseplate. Interacting residues are highlighted in orange. **I)** Tomograms of *Rhodobacter* phage RcSimone-Håstad showing a thin RcGTA-type baseplate (red arrows). Scale bar is 100 nm. **J)** Comparison of new baseplate protein models with previously published models (6tea,6teb,6teh), the current model is shown in ribbon and coloured based on the RMSD_C_α. Newly built regions are shown in cyan. Numbers represent delimiting residues.

To gain further insight into the molecular mechanism of TspA attachment to the baseplate, we optimized RcGTA purification conditions to obtain virions retaining an intact hexamer of TspA trimers (**Fig. S11**) and determined a high-resolution structure of the complex by single particle analysis, reaching an average resolution of 2.7 Å (**Fig. 5C, Fig. S12**). The 3D classification showed that TspA dissociates from the baseplate in pairs of trimers, as observed particles contained 0, 2, 4 or 6 adjacent trimers of TspA attached to the baseplate (**Fig. 5D**). A single TspA trimer is bound to the baseplate through a series of interactions involving: (i) the TspA C-terminal domain and the tail tube, (ii) TspA residues 205-320 and the distal tail protein OB-domain, and (iii) TspA loop 26-36 and the megatron peripheral domain loop 591-599 (**Fig. 5E-G, Table S3**). In virions lacking TspA, the tail tube loop 14-22, and the OB domain of distal tail protein, both involved in the TspA binding, have distorted density, suggesting their flexibility (19). We therefore conclude that the interaction with TspA stabilises their position and/or conformation.

The interface between two trimers of TspA has a buried area of 1,101 Å and a Δ^i^G of -5.5 kcal/mol (**Fig. 5H**). However, the overall buried surface area and calculated Δ^i^G of the TspA trimer-baseplate interface are 1,705 Å and -1.5 kcal/mol, respectively. In comparison, buried surface areas between other baseplate proteins, which range between 368 and 922 Å, and have much lower values of Δ^i^G, ranging between -5.6 and -11.7 kcal/mol per subunit (**Table S3**). Likewise, Δ^i^G is lower for the head spike-capsid assembly, with -4.5 kcal/mol and, based on a predicted model (**Data S3**), for the tail fibre-megatron assembly, with -13.7 kcal/mol (**Table S3**). The relatively high Δ^i^G value for the TspA-baseplate interface thus explains why TspA readily dissociate from the baseplate during purification and under *in situ* conditions.

Some tailed bacteriophages infecting *Rhodobacterales* share a homologous baseplate module with RcGTAs (36, 37), encoding all homologs that interact with TspA in the RcGTA virion, including the distal tail insertion domain (**Table S4**). However, PSI-BLAST searches did not identify any TspA-like homologs in phage sequences, apart from a 46-71 residues-long region in the cap domain (**Fig. S5**). We therefore tested whether phages instead harness the GTA component encoded by the host. For this, we imaged a crude lysate of a representative RcGTA-like baseplate phage, RcSimone-Håstad (36). CryoET reconstruction showed that in tested conditions, its baseplate lacks a TspA-like complex (**Fig. 5I**).

The resolution of the RcGTA baseplate assembly was sufficient for *de novo* model building of the megatron peripheral and fibre-binding domains, as well as the distal tail protein insertion domain (**Fig. 5J**). Moreover, it enabled the reassessment of the identity of three α-helices located in the central region of the RcGTA baseplate, previously suggested to be formed by the hydrophobic N-termini of the megatron protein (19). Owing to the improved density of this region, combined with improved protein sequence identification algorithms (38), we found that the density instead corresponds to the C-termini of the tape measure protein (**Fig. 5J**, **Fig. S13**). This interpretation is consistent with recent observations in other noncontractile long-tailed phages (39, 40).

## Discussion

Bacteriophages possess several enzymatically active proteins on their virion, typically required to degrade protective layers of the host cell envelope or to facilitate the penetration of the cell wall. While several tailspikes active against exopolysaccharides have been discovered (41), their function is tied to the baseplate itself and usually has evolved to assist the phage in depolymerizing the protective surface layer of its host to facilitate their infection. To our knowledge, phages do not encode soluble depolymerases dedicated to breaking down the biofilm matrix, likely because a fully soluble enzyme would be readily dispersed and diluted in the environment.

We discovered that *Rhodobacterales* GTAs, which are under different evolutionary pressure than bacteriophages, encode a biofilm depolymerase TspA. Unlike canonical phage tailspikes, this depolymerase is only loosely attached to the RcGTA baseplate. This allows precise delivery of the enzyme to the site where host cells are recognized by other virion proteins, and to dissociate from the virion to disrupt the nearby EPS (**Fig. 6A**). In this way, RcGTA solves the problem of deploying a soluble depolymerase which is still target-delivered to the correctly-recognised host.

**Fig. 6.**
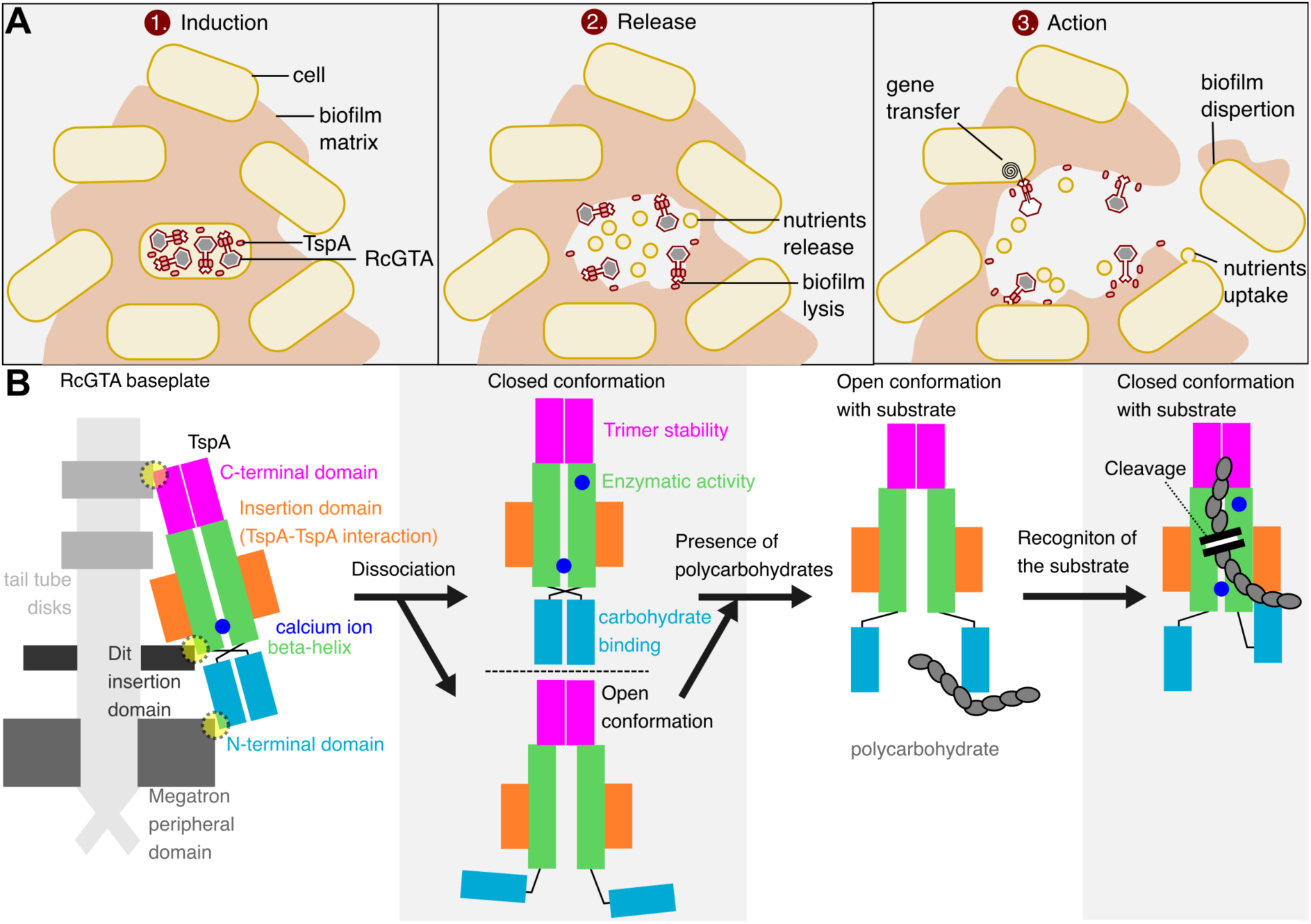
The proposed mechanisms of the TspA action. **A)** A scheme of events caused by RcGTA-TspA release. 1.) Particles are induced in biofilms in a subpopulation of cells. 2.) The cell lysis results in a release of nutrients, GTA virions with attached TspA and some soluble TspA. The nearby biofilm matrix is lysed by TspA activity. 3.) The nutrients are absorbed, genes transferred, and biofilm remodulated and potentially dispersed. **B)** A scheme of the molecular function of the TspA. Individual TspA domains are highlighted in different colours, as shown in the figure. The interaction interfaces between the TspA and the baseplate domains are highlighted in yellow circles. The calcium ions (blue circles) are essential for protein activity.

TspA has an N-terminal β-sandwich domain that becomes flexible upon detachment from the baseplate. This domain differs from the N-terminal baseplate-anchoring domain of phage tailspikes. We show that the β-sandwich domain shifts position upon substrate recognition. The flexibility of the domain likely serves as a metastable controller that dissociates the complex from the baseplate and allows for guiding the substrate into the active site. The active site of TspA is present in the interdomain conserved cleft, similarly as in the tailspike of coliphage Sf6 (42), and is stabilized by the coordination of calcium ions.

Interestingly, the major repressor of RcGTAs, the Rcc00280 protein, has been proposed to be an extracellular RTX family calcium-binding protein (27). This raises the possibility that calcium plays a broader role in controlling RcGTA production and activity.

Transcriptomic and proteomic analyses suggest that TspA is produced in quantities similar to those of the most abundant RcGTA component, the major capsid protein (**Fig. 1A, Table S1**). This raises the possibility that, in addition to the RcGTA virion-associated TspA, additional soluble TspA is released from the producer cell during lysis. Due to the small size of individual TspA trimers and the complex environment of the lysing cells, we were unable to directly confirm the presence of soluble TspA from the tomograms. However, this hypothetical soluble TspA would degrade the EPS that directly surrounds the producer cell, ensuring that RcGTA virion-associated TspA remains attached until it recognises the matrix near recipient cells.

The discovery of biofilm-degrading activity of TspA directly supports the hypothesis that the production of *Rhodobacterales* GTAs under native conditions occurs within biofilms and that the GTA-producing phenotype is selected for within biofilm subpopulations rather than among free-living planktonic cells (9). Overall, these findings show that RcGTA is a special phage-like entity shaped by evolution to function within biofilm matrix. TspA is a key component that enables the particle to reach EPS-embedded recipients. The identified molecular mechanism of TspA delivery and substrate recognition is key to inspiring the design of the next generation of biofilm depolymerases and phage-based strategies for eradicating industrial and medical biofilms.

## Acknowledgements

The research leading to these results has received funding from the Wellcome Trust grants 224067/Z/21/Z to P.B., 224665 to A.A.A., 109363/Z/15/A to P.C.M.F., and BBSRC grant UKRI2958 to P.C.M.F. Some of this work was performed at the Stanford-SLAC Cryo-EM Center (S2C2), which is supported by the National Institute of General Medical Sciences (1R24GM154186). The content is solely the responsibility of the authors and does not necessarily represent the official views of the National Institutes of Health. The authors would also like to thank to Prof. Wah Chiu (S2C2, Stanford-SLAC) for his invaluable support and assistance, Dr. Huw Jenkins (University of York) for discussions about single particle analysis, and Prof. Alan Davidson (University of Toronto) for mentoring P.B. Some of the imaging was done at the University of York cryo-EM facility supported by the Wellcome Trust (206161/Z/17/Z). The Viking cluster was used during this project, which is a high-performance compute facility provided by the University of York. We are grateful for computational support from the University of York, IT Services and the Research IT team. Further computing was performed on workstations purchased thanks to financial support from the Hartshorn-Jones fund at the University of York. Mass spectrometry analysis was performed at the York Centre of Excellence in Mass Spectrometry supported by a major capital investment through Science City York and Yorkshire Forward with funds from the Northern Way Initiative and EPSRC (EP/K039660/1; EP/M028127/1).

## Methods

### Growth of bacterial strains

Unless otherwise stated, the propagation of *Rhodobacter* strains was performed in rich media YPS, *E. coli* strains were grown in LB, with *Rhodobacter* strains incubated aerobically at 30°C and *E. coli* strains at 37°C.

### Quantitative PCR

Triplicate colonies of SB1003, SB1003:Δ*gafA*(gentR), SB1003:Δ*280*(specR) and SB1003:Δ*280*Δ*gafA*(specR-gentR) were inoculated in 5mL of YPS containing 10 µg/ml of kanamycin, and incubated at 30°C, 200 rpm for 24 hours aerobically. Then, 0.2 ml of the cultures was sub-cultured into 13.8 ml of RCV containing 10 µg/ml of Kanamycin, and incubated for an additional 72 hours, anaerobically. Then, 1 ml of cells was taken for total RNA purification using the NucleoSpin RNA kit (Macherey-Nagel) and treated with DNaseI on the column. The concentration of RNA was measured using NanoDrop, and 0.5 μg of total RNA was taken for PrimeScript FAST RT (Takara Bio) gDNA removal and reverse-transcription into cDNA template. One μl of 1:2 diluted cDNA template was used for each qPCR reaction with Fast SYBR Green Master-mix (Applied Biosystems) under standard conditions and 60°C annealing temperature (all efficiencies between 90-110%) using a QuantStudio 3 Real-Time PCR system. ΔCt values were calculated via Ct (gene of interest) – Ct (HKG (uvrD)), and used for statistical significance calculations of the samples. ΔΔCt values were calculated via ΔCt – Calibrator (average Δct of control group), and 2^-(ΔΔct) were used to represent log2 transformations of relative gene expression changes.

### Electromobility shift assay

For all DNA binding substrates, 50 base 5’-Cy5-labelled oligonucleotides (IDT) were annealed to complementary unlabelled oligonucleotides (IDT). Both oligos were mixed to a final concentration of 40 µM in annealing buffer (100 mM Potassium Acetate, 30 mM HEPES; pH 7.5) and heated to 98°C for 5 min then allowed to cool to room temperature.

Final concentrations in ten microliter EMSA mixtures were: 8 nM annealed Cy5-dsDNA, standard binding buffer (0.5X PBS, pH 8.0; 5 mM CaCl_2_; 5 mM MgCl_2_; 0.05% w/v Tween-20; 0.5 mg/ml BSA), 0.5 µg poly dI:dC, 4% glycerol and the specified concentrations of purified protein. Five-hundred-fold excess of competitor DNA was added to control mixtures. The specific competitor was unlabelled but otherwise identical to the binding substrate and the non-specific competitor was an unlabelled 50 bp annealed oligo from the *R. capsulatus* P*_ctrA_* promoter. All assays were incubated for 30 min at room temperature then immediately loaded onto a 7% Acrylamide gel (1 x TBE). Gels were run at 80 V for 90 min at room temperature in 1 x TBE. Fluorescence was imaged using a Typhoon Biomolecular Imager (Amersham) and analysed using ImageQuant (Amersham) and FIJI (43) software.

### Cloning and heterologous expression of TspA variants

The genes were amplified using Phusion polymerase from the colonies of *R. capsulatus* SB1003, *Tritonibacter mobilis* DSM23403 and *Phaeobacter piscinae* S26, respectively, utilising primers listed in **Table S5**. The amplified products were then cloned into linearized pETTFPP1 (pETYSBLIC3c) (44) using InPhusion (Takara) and transformed into *E. coli* Stellar cells (Takara). The correct sequence was verified using sequencing (Plasmidosaurus). The point mutations in TspA were generated in plasmid pETTFPP1_TspA by EMILI (45), using primers listed in **Table S5**. The mutated plasmids were transformed into *E. coli* Stellar cells and the presence of the mutation was verified by sequencing (Plasmidosaurus). Afterwards, all the constructs were transformed into *E. coli* BL21(DE3) gold. The cells were grown in 30°C/180 rpm to OD_600_=0.5, induced with 0.4 mM IPTG and expressed at 20°C/180 rpm overnight.

### Purification of TspA variants from *E. coli*

The cells were harvested, resuspended in the equilibration buffer (Tris 20mM, pH=7.8; NaCl 500mM, imidazole 20mM), and sonicated using Sonoplus HD2070 (Bandelin). The lysate was spun at 24,000 G/20 min and loaded onto the HisTrap 5ml FF column (Cytiva). The protein was eluted using a linear gradient of elution buffer (Tris 20mM, pH=7.8; NaCl 500mM, Imidazole 500mM). The fractions corresponding to the peak were collected, concentrated and buffer exchanged to the equilibration buffer using 30 kDa Amicon® Ultra filter units (Merck Millipore). The His-tag was cleaved using 3C protease (prepared by YSBL, University of York) in a mass ratio of protease to protein equal to 1:100, by incubating the mixture at 4°C overnight. The sample was then loaded onto equilibrated Super Ni-NTA resin (Generon), incubated for 20 min, and spun at 3,000 G/2 min. The unbound protein was loaded onto Superose 6 increase 10/300 GL, and the peak corresponding to the protein trimer (elution volume ∼ 15.5 ml) was fractionated and concentrated using the filter units. The sample concentration was estimated using A_280_ readings, and the protein was flash-frozen and kept at -70°C.

### Purification of TspA from *Rhodobacter*

*R. capsulatus* strains SB1003Δ*rcc00280* and SB1003Δ*rcc00280*Δ*tspA* were individually inoculated into RCV media, grown at 30°C, shaking at 190 rpm until the stationary phase, and inoculated 1:100 into fresh YPS media in 25 ml screw cap tubes. The samples were incubated at 30°C in an illumination cabinet, placed 30 cm from three 40 W light-emitting tubes (Panasonic FL40SS・W/37c). After the cultivation, samples were centrifuged at 10,000 G/30 min/4°C, and the supernatant was carefully taken and consecutively filtered through 0.8 and 0.4 µm Supor^TM^ PES filters (Pall Life Sciences). This supernatant was then diluted 1:2 with a loading buffer (50 mM K-phosphate, pH 7.0; 10 mM NaCl; 5 mM MgSO_4_) and run through a CIM-multus QA-HR 1ml column (Sartorius BIASeparations) using a step gradient, with 4 % elution buffer (conductivity of ∼ 12 mS/cm) used for eluting the particles. The corresponding fractions were pooled and concentrated using 30kDa PES filter spin columns (VivaSpin) and run through Superose 6 increase 10/300 GL (Cytiva) equilibrated with 50 mM K-phosphate, pH 7.0; 90mM NaCl, 5 mM MgSO_4_. The final sample was further concentrated using the filter spin column to A_280_=5.9, flash frozen and kept at -80°C.

### Biofilm assays

*R. capsulatus* SB1003 and *C. sphaeroides* 2.4.1. were inoculated into YPS and grown aerobically overnight. The culture was diluted 1:100 to fresh YPS media, and 200 µl were added per single well to a 96-well microplate (Corning). In the case of *R. mobilis* and *P. piscinae*, Marine Broth (Difco) was used instead of the YPS. Then, for the biofilm prophylaxis assay, 12 µg of TspA protein or buffer of the same volume were added per well and mixed. The plates were wrapped in parafilm to restrain evaporation and incubated at 25°C (in case of *Phaeobacter* TspA) or 30°C (in case of *Rhodobacter* TspA) for 72 hours. The wells were washed 3 times in water, then stained with 0.1 % crystal violet for 20 min, again washed 3 times with water, resuspended in 96% ethanol, and the OD_595_ was measured using a Tecan plate reader. In the case of biofilm disposal assay, the plates were incubated for 48 hours, wells were washed 3 times with water, then 200 µl of fresh media were added (YPS or YP salt derivatives), together with 12 µg of TspA protein or buffer of the same volume.

The plate was then incubated for 2 hours at 30°C, wells were washed 3 times with water, resuspended in 96% ethanol, and the OD_595_ was measured using the Tecan plate reader. In case of pre-treatment with catalytically-dead variants, the well was first treated with 12 µg of TspA variant or buffer for 2 hours at 30°C, then 12 µg wildtype TspA were added, and the incubation continued for another 2 hours.

### Gene transfer assays

*Rhodobacter* assays were carried out essentially as described in Leung and Beatty, 2013 (46). RcGTA donor cultures were grown under anoxic conditions with illumination in YPS for 72 h. Cells were cleared from donor cultures by centrifugation and the supernatant filtered through a 0.45 µm syringe filter. Recipient cultures were grown either under chemotrophic conditions in RCV for 24 h or anoxic illuminated conditions for 72 h, as indicated in the text. Recipient cells were concentrated 3-fold by centrifugation at 5,000 x g and resuspension in 1/3 volume of G-Buffer (10 mM Tris-HCl, pH 7.8; 1 mM MgCl_2_; 1 mM CaCl_2_; 1 mM NaCl; 0.5 mg/ml BSA). Where stated, purified TspA protein was added to the recipient cells and incubated for 5 min at room temperature. Reactions were carried out in polystyrene culture tubes (Starlab) containing 400 µl G-Buffer, 100 µl recipient cells and 100 µl filter donor supernatant, and incubated at 30°C for 1 h. Nine-hundred microlitres of YPS was added to each tube and incubated for a further 3 h. Cells were harvested by centrifugation at 5,000 x g and plated on YPS plates containing 100 µg/ml rifampicin.

### TspA sample preparation and cryo-EM data acquisition

The purified TspA was diluted to a final concentration of 0.25 mg/ml in buffer (20mM Tris, pH=7.5; NaCl 250mM) or in YPS, and 3.8 µl of the sample was applied onto Quantifoil R3.5/1 300 mesh copper grids coated with 2nm carbon, and vitrified using Vitrobot IV (Thermo Fisher Scientific). The vitrification parameters were 100 % humidity, 0 force, 8 s wait time and 2 s blotting time. The grids were screened, and data acquired as stated in **Table S6**.

### Cryo-EM samples of TspA with a polysaccharide ligand

The purified TspA was diluted to a final concentration of 0.125 mg/ml in samples of extracellular polysaccharides produced by B10 and SB1003 strains. For B10 strains, 1 ml of a culture that was grown anaerobically in RCV for three days was spun at 1,500 G/5 min, the supernatant was transferred to a fresh tube and spun at 8,000 G/5 min. The pellet was then washed with 1 ml of fresh RCV three times, with each wash followed by 8,000 G/5 min spin. The final pellet was resuspended in 2 ml of YPS. For the SB1003 strain, the culture was grown in YPS on polystyrene plates with a flat well for 48 hours. Then, the wells were washed with water, and the firmly attached substrate at the bottom of 10 wells was thoroughly resuspended in 300 µl of YPS using the pipette tip to mechanically disrupt the substrate. The sample was then spun at 8,000 G/5 min, and the pellet was resuspended in 50 µl of fresh YPS. The grids were prepared, and data were collected under the same conditions as those used for pure TspA (**Table S6**).

### Purification of RcGTAs with intact TspA

*R. capsulatus* strain DE442 was incubated in the same way as reported for the purification of TspA. The filtrated supernatants were then diluted 1:2 with a loading buffer (20mM HEPES, pH 7.0; 10 mM NaCl; 5 mM CaCl_2_) and run through a CIM-multus QA-HR 1ml column (Sartorius BIASeparations) using a step gradient, with 11 % of elution buffer (20mM HEPES, pH 7.0; 1.5 M NaCl; 5 mM CaCl_2_) used for eluting the particles, which equaled to conductivity of ∼22 mS/cm (**Fig. S11**). The corresponding fractions were pooled and concentrated using 100kDa PES filter spin columns (VivaSpin), buffer transferred to modified G-buffer (10 mM Tris, pH 7.8; 1 mM NaCl, 1 mM MgCl_2_, 2.5 mM CaCl_2_, BSA 10 ug/ml) and ultracentrifuged in a 15-40% sucrose gradient based on the previous protocol (19). The bottom 0.5 ml fraction corresponding to RcGTAs was taken using a syringe with a cannula needle, buffer exchanged using modified G-buffer to contain <0.05% sucrose and concentrated to A_280_=30 mg/ml using the filter spin columns.

### RcGTA *in vitro* sample preparation and cryo-EM data acquisition

Four microliters of purified RcGTAs in a concentration of A_280_=30 mg/ml were applied onto glow-discharged Quantifoil R2/1 300 mesh copper grids and vitrified using Vitrobot IV (Thermo Fisher Scientific). The vitrification parameters were 100 % humidity, 0 force, 8 s wait time and 2 s blotting time. The grids were screened, and data acquired as stated in **Table S6.**

### Single particle analysis and model refinement

The analysis of purified TspA incubated in rich media YPS, buffer and with ligand, respectively, was processed in RELION5 (47), with the pipelines described in **Fig. S3**, **S6** and **S9**. The analysis of RcGTAs associated with TspA was performed in RELION5, with the pipeline described in **Fig. S12**. The final densities from local resolution runs were used for building the protein models. Here, a predicted structure by AlphaFold3 (48) was rigid body-fitted into the map using ChimeraX 1.10 (49) fitmap command and further manually refined in Coot 0.9.8.5 (50). Then, the structure was automatically refined in Phenix 1.20 (51) using real space refinement. After several rounds of manual coot and phenix real space refinement, the model was further refined in ChimeraX using ISOLDE pluggin (52), gradually refining 15 residue-long segments. The model was then refined in Phenix 1.20 using real space refinement and the waters were added and refined using a Phenix command phenix.douse, with H-bind distances to consider in range of 2.0-3.3 Å, and manually optimising the sphericity parameter, the other parameters were left default. The density was then checked for any missing waters with obvious density, which were added manually and refined in Coot and ISOLDE, with the final model refined in Phenix. The Q-score (53) was determined using ChimeraX plugin.

### RcGTA *in situ* sample preparation, cryo-ET data acquisition and pre-processing

*Rhodobacter capsulatus* SB1003 containing plasmid pQF with RcGTA transcriptional regulator gafA under the control of a cumate-inducible promoter was cultivated aerobically at 30°C, 200 rpm in RCV until the stationary phase. Then, cumate was added to the final concentration of 25 µM, and the incubation continued for another 3 hours. The cells were then mixed with 6 nm gold beads (Aurion) in a ratio of 8:1, and 3.8 ul of the mixture was applied onto glow-discharged 200 mesh R1.2/1.3 Quantifoil® carbon grids, and vitrified using Vitrobot IV (Thermo Fisher Scientific). The vitrification parameters were 70 % humidity, 0 force, 2 s wait time and 10 s blotting time. The front blotting pad was covered with parafilm, with the effective blotting occurring only with the back pad.

The grids were screened at the York cryo-EM facility using Glacios TEM (Thermo Fisher Scientific) operated at 200kV, and data were acquired as stated in **Table S6.** Raw movies were motion-corrected using MotionCor2 implementation in RELION (54), and tilt series stacks were generated and dose-weighted using the alignframes command in IMOD (55). The stacks were imported to EMAN2 (56), where final tomograms were generated and the subtomogram averaging performed.

### Subtomogram averaging of RcGTA *in situ*

Particles were picked by a reference-based picking using EMD-10592 as a reference. Picks were manually screened and bad hits removed. Final count of particles was 455. EMD-10592 low-passed to 60 Å was used as an initial model for subtomogram averaging, which resulted in a 30 Å map showing extra density of the baseplate region. The particles were then repicked using EMD-10565 as a reference, and any obvious left-out particles were added using manual picking, resulting in 902 particles. The subtomogram averaging was repeated, and the particles were subsequently classified into two classes without alignment using a mask encompassing the baseplate density. Bad particles were removed, leaving 551 good particles. The particles were then refined using local searches and a soft mask around the particle, resulting in a 21.1 Å map (**Fig. S10**).

Subparticles containing baseplates were extracted using coordinate 0,0,112 in a box size of 104. Metadata from C1 reconstruction were gathered, and particles refined using local searches with C3 symmetry. This led to a reconstruction with pseudo-C6 symmetry, as the signal from the hexameric region of the baseplate dominated the reconstruction. The particles were classified into 2 groups using a skipalign option and a mask covering only the bottom baseplate region containing tail fibres, which has a distinct c3 symmetry. This resulted in a separation of two trimeric densities rotated 60 degrees relative to each other. Particles from class 0 were retained, while those in class 1 were symmetry expanded using e2proclst.py in the C6 group, resulting in particle list symmetry copies. The group ID=1 was selected and merged with the list of particles from the original class 0 using e2proclst.py, yielding a complete list of particles, both oriented in the same C3 position. This list of particles was then refined with local searches and C3 symmetry, resulting in a final 14.7 Å map (**Fig. S10**).

### Propagation and imaging of phage RcSimone-Håstad

A single plaque of the phage was resuspended in 200 ul of YPS, and 10 µl of the suspension was dropped onto a log-phase culture of SB1003 plated on YPS plates overlayed with YPS 0.7% agar. After 1 day of incubation at 30°C and an additional day at 22°C, turbid areas corresponding to drops appeared. The soft agar of these areas was scooped using an inoculation loop, resuspended in 2 ml, and debris was settled for 30 min. Four µl from the top of the tube were applied onto glow-discharged Quantifoil 200 mesh R1.3/1.2 grid, back-blotted for 3.5 seconds at 65% humidity and vitrified using Vitrobot IV. Tilt series of the sample were acquired at ×45 000 magnification and -5 μm defocus using -21°,54° and -24°,-54° bidirectional tilt series acquisition scheme with 3° increment and a total dose of 3.2 e-.Å-2 per each tilt image. The tilt images were processed using automatic batch alignment and reconstructed using a SIRT-like filter with 15 iterations in IMOD 4.12.49 Tomography package Etomo (57).

### Bioinformatic analysis

The taxid analysis was performed using a PSI-BLAST search (58), limiting the results to hits with more than 75% query coverage (**Data S1**). The TspA conservation analysis was performed manually by blast searches and subsequent inspection of the genome location near *lysS* gene in Artemis (59). The sequence and fold similarity search was performed using blastp (58), HHpred (60) and FoldSeek (61).

## Data visualization

The structural figures were created using UCSF ChimeraX 1.11 (49). The tomograms were visualized using IMOD 4.12 (57). The charts were plotted using the ggplot2 system of the package R (https://www.r-project.org/).

## Statistical analysis

Statistical analysis was carried out using Excel and for each use, the test parameters are indicated in the figure legends. For the t-test, the function T.TEST was applied. For the statistical significance calculations of the samples in RT-qPCR, one-way ANOVA with Tukey Post-Hoc pairwise multiple comparisons were performed in Statistical Package for the Social Sciences (IBM). For the biofilm assays, the function ANOVA within the Analysis ToolPak add-in was used, and the Tukey-Kramer Post Hoc Test was performed using a formula “Q_critical value_ = Q × √(s^2^_pooled_ / n)”. Here, Q is the value from studentized range Q table (62), s^2^_pooled_ is pooled variance across all groups, n is sample size for a given group. The significance was then estimated by comparing the absolute mean difference for each group with the Q_critical value_ (**Data S2**).

## References

1. M. W. Craske, J. S. Wilson, P. C. M. Fogg, Gene transfer agents: structural and functional properties of domesticated viruses. Trends in Microbiology (2024). 10.1016/j.tim.2024.05.002.

2. M. N. Price, et al., Mutant phenotypes for thousands of bacterial genes of unknown function. Nature 557, 503–509 (2018).

3. R. Kogay, et al., Machine-Learning Classification Suggests That Many Alphaproteobacterial Prophages May Instead Be Gene Transfer Agents. Genome Biol Evol 11, 2941–2953 (2019).

4. K. Gozzi, N. T. Tran, J. W. Modell, T. B. K. Le, M. T. Laub, Prophage-like gene transfer agents promote Caulobacter crescentus survival and DNA repair during stationary phase. PLOS Biology 20, e3001790 (2022).

5. L. D. McDaniel, et al., High frequency of horizontal gene transfer in the oceans. Science 330, 50 (2010).

6. L. D. McDaniel, E. C. Young, K. B. Ritchie, J. H. Paul, Environmental factors influencing gene transfer agent (GTA) mediated transduction in the subtropical ocean. PLoS One 7, e43506 (2012).

7. P. C. M. Fogg, A. B. Westbye, J. T. Beatty, One for All or All for One: Heterogeneous Expression and Host Cell Lysis Are Key to Gene Transfer Agent Activity in Rhodobacter capsulatus. PLOS ONE 7, e43772 (2012).

8. S. Koppenhöfer, et al., Integrated Transcriptional Regulatory Network of Quorum Sensing, Replication Control, and SOS Response in Dinoroseobacter shibae. Front. Microbiol. 10 (2019).

9. R. Kogay, O. Zhaxybayeva, Co-evolution of gene transfer agents and their alphaproteobacterial hosts. Journal of Bacteriology 206, e00398–23 (2024).

10. R. J. Redfield, S. M. Soucy, Evolution of Bacterial Gene Transfer Agents. Front Microbiol 9, 2527 (2018).

11. P. C. M. Fogg, Gene transfer agents: The ambiguous role of selfless viruses in genetic exchange and bacterial evolution. Mol Microbiol 123, 124–131 (2025).

12. L. Šmerdová, et al., Dynamics of bacterial biofilm development imaged using light sheet fluorescence microscopy. Biochem Biophys Rep 43, 102127 (2025).

13. D. Sherlock, P. C. M. Fogg, Loss of the Rhodobacter capsulatus Serine Acetyl Transferase Gene, cysE1, Impairs Gene Transfer by Gene Transfer Agents and Biofilm Phenotypes. Appl Environ Microbiol 88, e0094422 (2022).

14. M. Valentini, A. Filloux, Biofilms and Cyclic di-GMP (c-di-GMP) Signaling: Lessons from Pseudomonas aeruginosa and Other Bacteria. The Journal of Biological Chemistry 291, 12547 (2016).

15. P. Pallegar, L. Peña-Castillo, E. Langille, M. Gomelsky, A. S. Lang, Cyclic di-GMP-Mediated Regulation of Gene Transfer and Motility in Rhodobacter capsulatus. Journal of Bacteriology 202, e00554 (2020).

16. S. Koppenhöfer, A. S. Lang, Interactions among Redox Regulators and the CtrA Phosphorelay in Dinoroseobacter shibae and Rhodobacter capsulatus. Microorganisms 8, 562 (2020).

17. P. Pallegar, M. Canuti, E. Langille, L. Peña-Castillo, A. S. Lang, A Two-Component System Acquired by Horizontal Gene Transfer Modulates Gene Transfer and Motility via Cyclic Dimeric GMP. Journal of Molecular Biology 432, 4840–4855 (2020).

18. L. L. Lindqvist, et al., Tropodithietic Acid, a Multifunctional Antimicrobial, Facilitates Adaption and Colonization of the Producer, Phaeobacter piscinae. mSphere 8, e0051722 (2023).

19. P. Bárdy, et al., Structure and mechanism of DNA delivery of a gene transfer agent. Nat Commun 11, 3034 (2020).

20. A. P. Hynes, et al., Functional and Evolutionary Characterization of a Gene Transfer Agent’s Multilocus “Genome.” Molecular Biology and Evolution 33, 2530–2543 (2016).

21. T. B. Stanton, S. B. Humphrey, D. O. Bayles, R. L. Zuerner, Identification of a divided genome for VSH-1, the prophage-like gene transfer agent of Brachyspira hyodysenteriae. J Bacteriol 191, 1719–1721 (2009).

22. R. Kogay, O. Zhaxybayeva, Selection for Translational Efficiency in Genes Associated with Alphaproteobacterial Gene Transfer Agents. mSystems 7, e00892–22 (2022).

23. R. Kogay, Y. I. Wolf, E. V. Koonin, O. Zhaxybayeva, Selection for Reducing Energy Cost of Protein Production Drives the GC Content and Amino Acid Composition Bias in Gene Transfer Agents. mBio 11, 10.1128/mbio.01206-20 (2020).

24. A. B. Westbye, K. Kuchinski, C. K. Yip, J. T. Beatty, The Gene Transfer Agent RcGTA Contains Head Spikes Needed for Binding to the Rhodobacter capsulatus Polysaccharide Cell Capsule. Journal of Molecular Biology 428, 477–491 (2016).

25. P. C. M. Fogg, Identification and characterization of a direct activator of a gene transfer agent. Nat Commun 10, 595 (2019).

26. P. Bardy, et al., Penton blooming, a conserved mechanism of genome delivery used by disparate microviruses. mBio 16, e0371324 (2025).

27. H. Ding, M. P. Grüll, M. E. Mulligan, A. S. Lang, J. T. Beatty, Induction of Rhodobacter capsulatus Gene Transfer Agent Gene Expression Is a Bistable Stochastic Process Repressed by an Extracellular Calcium-Binding RTX Protein Homologue. Journal of Bacteriology 201, 10.1128/jb.00430-19 (2019).

28. A. S. Omar, J. Weckesser, H. Mayer, Different polysaccharides in the external layers (capsule and slime) of the cell envelope of Rhodopseudomonas capsulata Sp11. Arch Microbiol 136, 291–296 (1983).

29. G. A. O’Toole, Microtiter Dish Biofilm Formation Assay. J Vis Exp 2437 (2011). 10.3791/2437.

30. A. A. Burnim, K. Dufault-Thompson, X. Jiang, The three-sided right-handed β-helix is a versatile fold for glycan interactions. Glycobiology 34, cwae037 (2024).

31. K. Stummeyer, A. Dickmanns, M. Mühlenhoff, R. Gerardy-Schahn, R. Ficner, Crystal structure of the polysialic acid–degrading endosialidase of bacteriophage K1F. Nat Struct Mol Biol 12, 90–96 (2005).

32. P. Youkharibache, et al., The Small β-Barrel Domain: A Survey-Based Structural Analysis. Structure 27, 6–26 (2019).

33. T. Itoh, et al., Crystal structure of the catalytic unit of thermostable GH87 α-1,3-glucanase from *Streptomyces thermodiastaticus* strain HF3-3. Biochemical and Biophysical Research Communications 533, 1170–1176 (2020).

34. E. Krissinel, K. Henrick, Inference of Macromolecular Assemblies from Crystalline State. Journal of Molecular Biology 372, 774–797 (2007).

35. J. Pei, N. V. Grishin, AL2CO: calculation of positional conservation in a protein sequence alignment. Bioinformatics 17, 700–712 (2001).

36. J. Rapala, et al., Genomic diversity of bacteriophages infecting Rhodobacter capsulatus and their relatedness to its gene transfer agent RcGTA. PLoS One 16, e0255262 (2021).

37. Y. Huang, et al., Structure and proposed DNA delivery mechanism of a marine roseophage. Nat Commun 14, 3609 (2023).

38. K. Jamali, et al., Automated model building and protein identification in cryo-EM maps. Nature 628, 450–457 (2024).

39. R. Linares, et al., Structural basis of bacteriophage T5 infection trigger and E. coli cell wall perforation. Science Advances 9, eade9674 (2023).

40. J. L. Kizziah, K. A. Manning, A. D. Dearborn, T. Dokland, Structure of the host cell recognition and penetration machinery of a Staphylococcus aureus bacteriophage. PLOS Pathogens 16, e1008314 (2020).

41. A. Visnapuu, M. Van der Gucht, J. Wagemans, R. Lavigne, Deconstructing the Phage–Bacterial Biofilm Interaction as a Basis to Establish New Antibiofilm Strategies. Viruses 14, 1057 (2022).

42. Y. Kang, et al., Bacteriophage Tailspikes and Bacterial O-Antigens as a Model System to Study Weak-Affinity Protein–Polysaccharide Interactions. J. Am. Chem. Soc. 138, 9109–9118 (2016).

43. J. Schindelin, et al., Fiji: an open-source platform for biological-image analysis. Nat Methods 9, 676–682 (2012).

44. M. J. Fogg, A. J. Wilkinson, Higher-throughput approaches to crystallization and crystal structure determination. Biochemical Society Transactions 36, 771–775 (2008).

45. T. Füzik, P. Ulbrich, T. Ruml, Efficient Mutagenesis Independent of Ligation (EMILI). J Microbiol Methods 106, 67–71 (2014).

46. M. M. Leung, J. T. Beatty, <EM>Rhodobacter capsulatus</EM> Gene Transfer Agent Transduction Assay. Bio-protocol 3 (2013).

47. D. Kimanius, L. Dong, G. Sharov, T. Nakane, S. H. W. Scheres, New tools for automated cryo-EM single-particle analysis in RELION-4.0. Biochem J 478, 4169–4185 (2021).

48. J. Abramson, et al., Accurate structure prediction of biomolecular interactions with AlphaFold 3. Nature 630, 493–500 (2024).

49. E. F. Pettersen, et al., UCSF ChimeraX: Structure visualization for researchers, educators, and developers. Protein Sci 30, 70–82 (2021).

50. P. Emsley, B. Lohkamp, W. G. Scott, K. Cowtan, Features and development of Coot. Acta Crystallogr D Biol Crystallogr 66, 486–501 (2010).

51. D. Liebschner, et al., Macromolecular structure determination using X-rays, neutrons and electrons: recent developments in Phenix. Acta Crystallogr D Struct Biol 75, 861–877 (2019).

52. T. I. Croll, ISOLDE: a physically realistic environment for model building into low-resolution electron-density maps. Acta Cryst D 74, 519–530 (2018).

53. G. Pintilie, et al., Q-score as a reliability measure for protein, nucleic acid and small-molecule atomic coordinate models derived from 3DEM maps. Acta Cryst D 81, 410–422 (2025).

54. J. Zivanov, et al., New tools for automated high-resolution cryo-EM structure determination in RELION-3. eLife 7, e42166 (2018).

55. J. R. Kremer, D. N. Mastronarde, J. R. McIntosh, Computer visualization of three-dimensional image data using IMOD. J Struct Biol 116, 71–76 (1996).

56. M. Chen, et al., A complete data processing workflow for cryo-ET and subtomogram averaging. Nat Methods 16, 1161–1168 (2019).

57. J. R. Kremer, D. N. Mastronarde, J. R. McIntosh, Computer visualization of three-dimensional image data using IMOD. J Struct Biol 116, 71–76 (1996).

58. S. F. Altschul, et al., Gapped BLAST and PSI-BLAST: a new generation of protein database search programs. Nucleic Acids Research 25, 3389–3402 (1997).

59. T. Carver, S. R. Harris, M. Berriman, J. Parkhill, J. A. McQuillan, Artemis: an integrated platform for visualization and analysis of high-throughput sequence-based experimental data. Bioinformatics 28, 464–469 (2012).

60. L. Zimmermann, et al., A Completely Reimplemented MPI Bioinformatics Toolkit with a New HHpred Server at its Core. J Mol Biol 430, 2237–2243 (2018).

61. M. van Kempen, et al., Fast and accurate protein structure search with Foldseek. Nat Biotechnol 42, 243–246 (2024).

62. H. L. Harter, Tables of Range and Studentized Range. The Annals of Mathematical Statistics 31, 1122–1147 (1960).

