## Supplementary Information for "The mechanism of biofilm degradation by a detachable tailspike of gene transfer agents"

Pavol Bardy *et al.*

**This PDF file includes:**

Figs. S1 to S13
Tables S1 to S6
Supplementary references 1 to 11

**Other Supplementary Materials for this manuscript include the following:**
Data S1 to S3

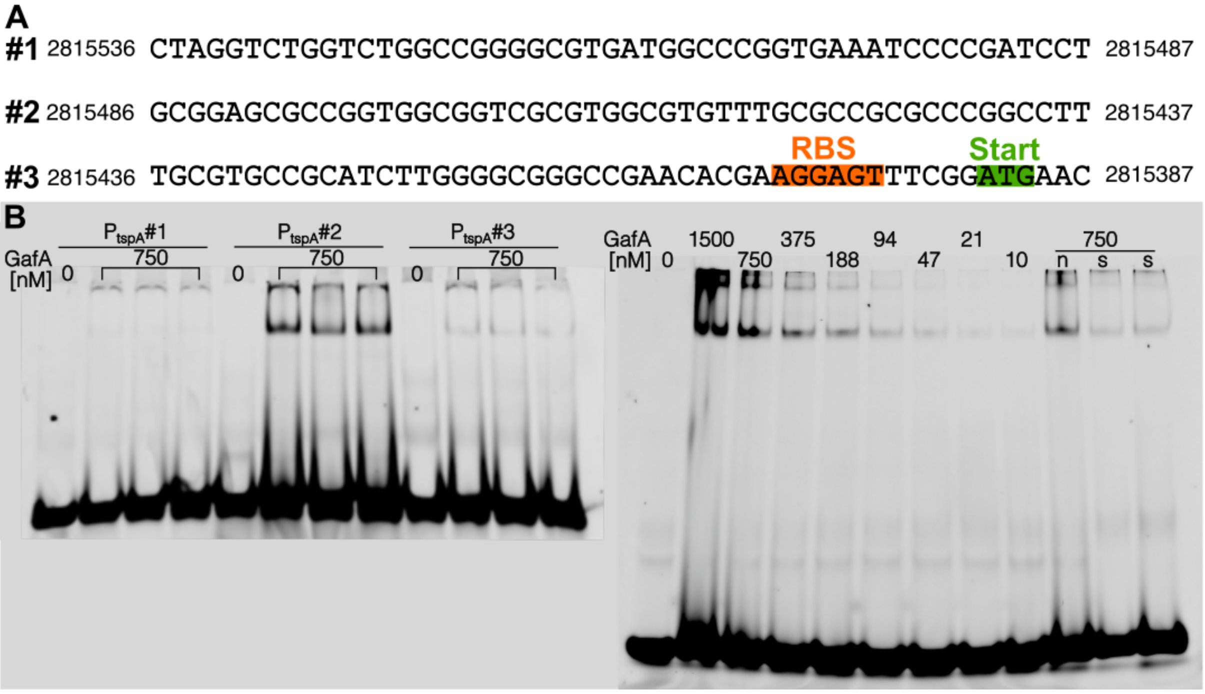


**Fig. S1. Data related to the electromobility shift assay. A)** Sequence of the regions upstream of *tspA,* which were used for the electromobility shift assay with GafA. The complete region corresponds to the 2815387-2815536 of GenBank entry CP001312.1**.** RBS, ribosome binding site of *tspA*; Start, start codon of *tspA*. **B)** Uncropped gels, cropped versions are shown in the main text, **Fig. 1**.

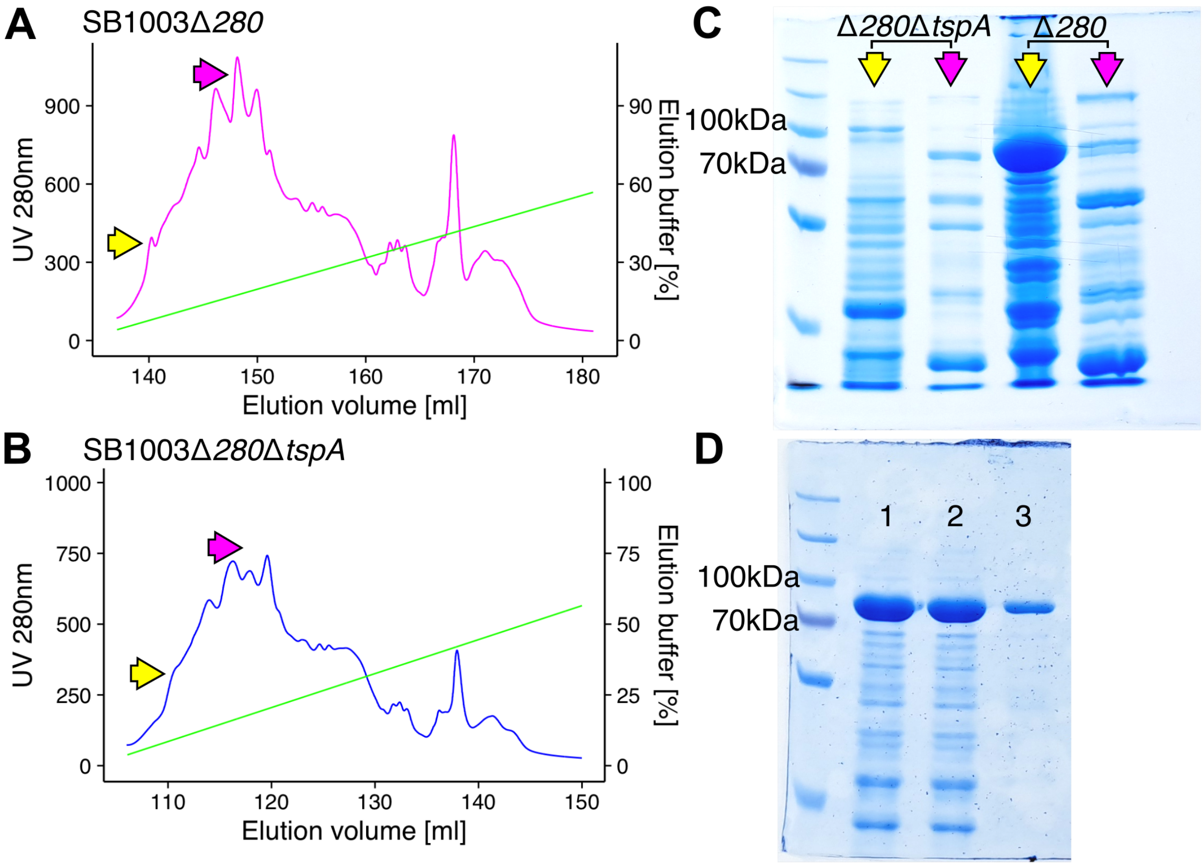


**Fig. S2. Purification of TspA from *R. capsulatus*. A-B)** Chromatograms of supernatants from SB1003Δ*280* (A) and SB1003Δ*280*Δ*tspA* (B) run through an ion-exchange CIMmultus ® monolith QA-HR column. Two peaks, which significantly differed between the two samples, were identified and are highlighted by arrows. **C)** SDS-PAGE of the peaks shown in panels A and B. The expected size of TspA is 81.7kDa, corresponding to the strong band observed in the peak highlighted by the yellow arrow in the Δ*280* strain. **D)** Further purification of TspA via size exclusion. 1, pooled sample after ion-exchange chromatography; 2, sample after a further buffer-exchange; 3, pure sample after the size-exclusion. The marker is PageRuler Plus Prestained Protein Ladder (10-250kDa, ThermoFisher Scientific).

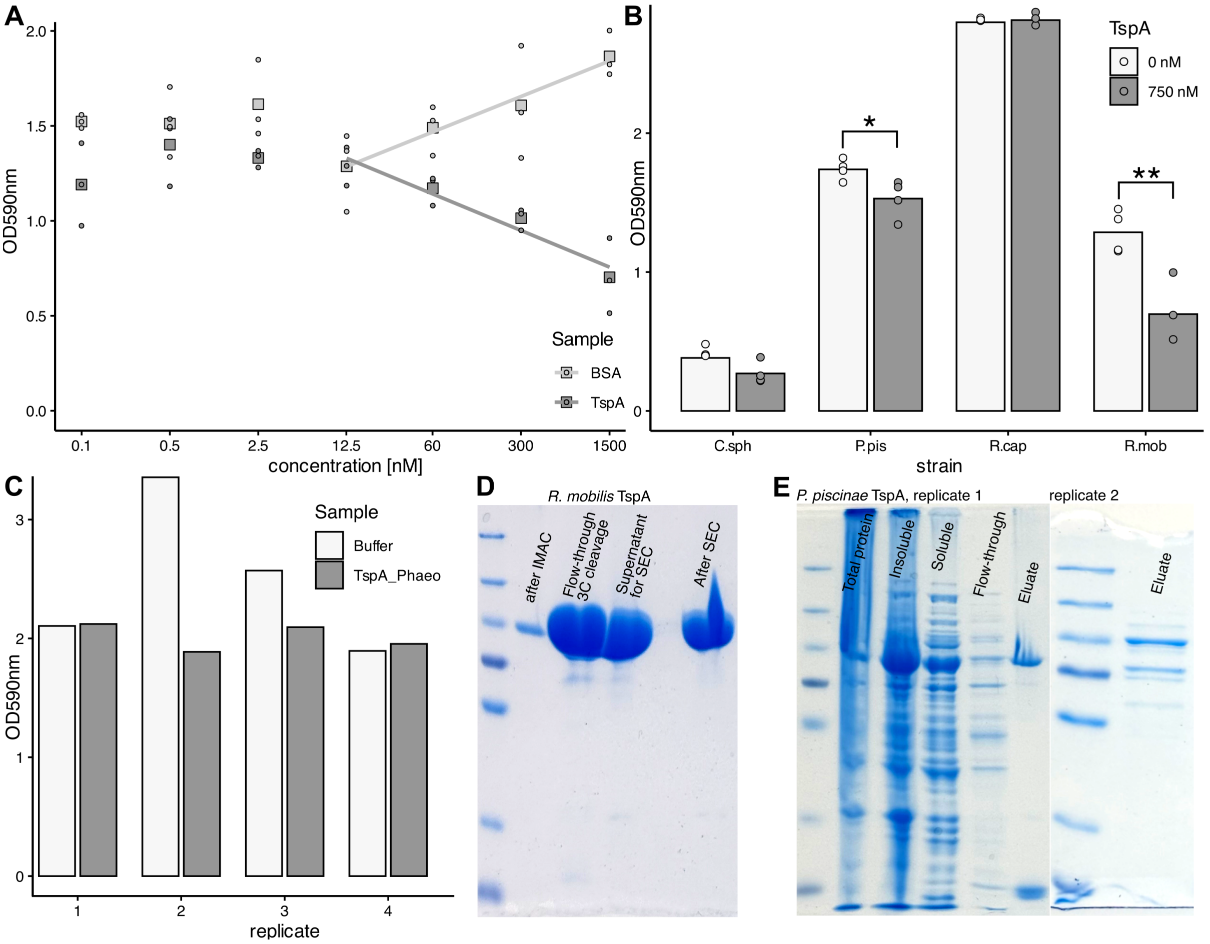


**Fig. S3. Effect of TspA homologs from *Ruegeria mobilis* on biofilm removal. A)** The level of biofilm formation by *R. mobilis* DSM15170 upon 72-hour incubation after addition of different concentrations of buffer or *R. mobilis* homolog TspA at time zero. A trend line is shown for concentrations 12.5 - 1500 nM. **B)** The level of biofilm formation by different *Rhodobacterales* species upon 72-hour incubation after addition of buffer or *R. mobilis* homolog of TspA at time zero. Csph, *Cereibacter sphaeroides*; Ppis, *Phaeobacter piscinae*; Rcap, *Rhodobacter capsulatus*; Rmob, *Ruegeria mobilis*. Statistical significance was calculated via paired t-test (**, $p<0.005$; *, $p <0.05$; and $p >0.05$, not shown). **C)** The level of biofilm formation by *P. piscinae* upon 72-hour incubation after addition of buffer or crudely-purified *P. piscnae* homolog of TspA at time zero. Results of four independent expressions, purifications and biofilm assays are shown. **D)** SDS-PAGE of purified TspA homolog of *R. mobilis.* IMAC, immobilized metal-affinity chromatography; SEC, size-exclusion chromatography. **E)** SDS-PAGE of crudely purified TspA homolog of *P. piscinae*. The eluate from Ni-NTA agarose beads of two replicates is shown. The marker in panels D-E) is PageRuler Plus Prestained Protein Ladder (10-250kDa, ThermoFisher Scientific).

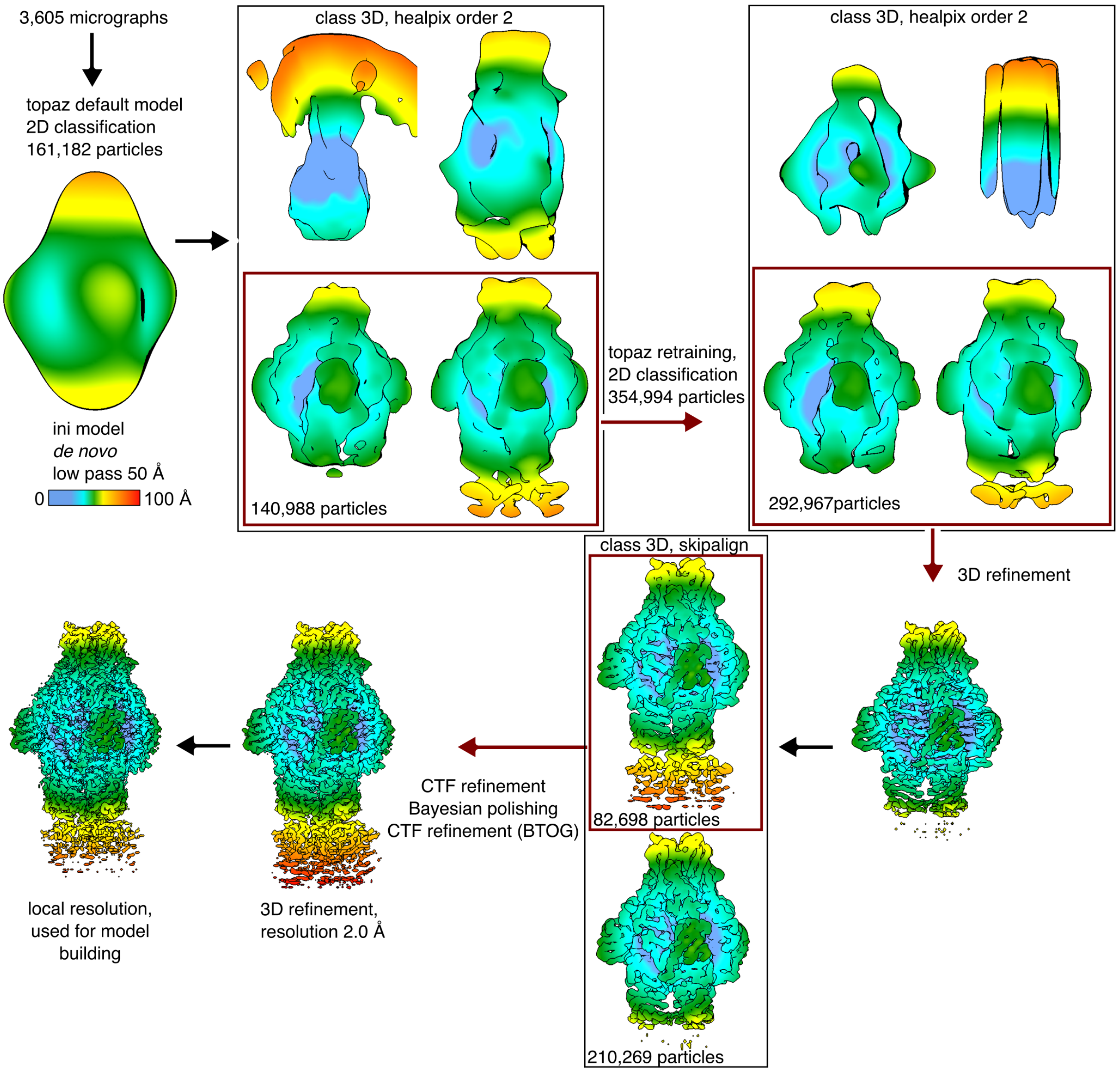


**Fig. S4. Single particle analysis of TspA incubated in YPS.** The analysis was done in RELION5 (*1*). The colour bar indicates the distance from the centre of the particle. The number of micrographs corresponds to the list filtered according to defocus, astigmatism and estimated resolution. The initial model was taken from the dataset of TspA incubated in buffer, where it was calculated *de novo*. Another 3D refinement was run after CTF refinement and Bayesian polishing. BTOG, beam tilt optics group, with the star file optics group edited using starpy scripts made by Dr Tibor Fuzik, CEITEC (https://github.com/fuzikt/starpy).

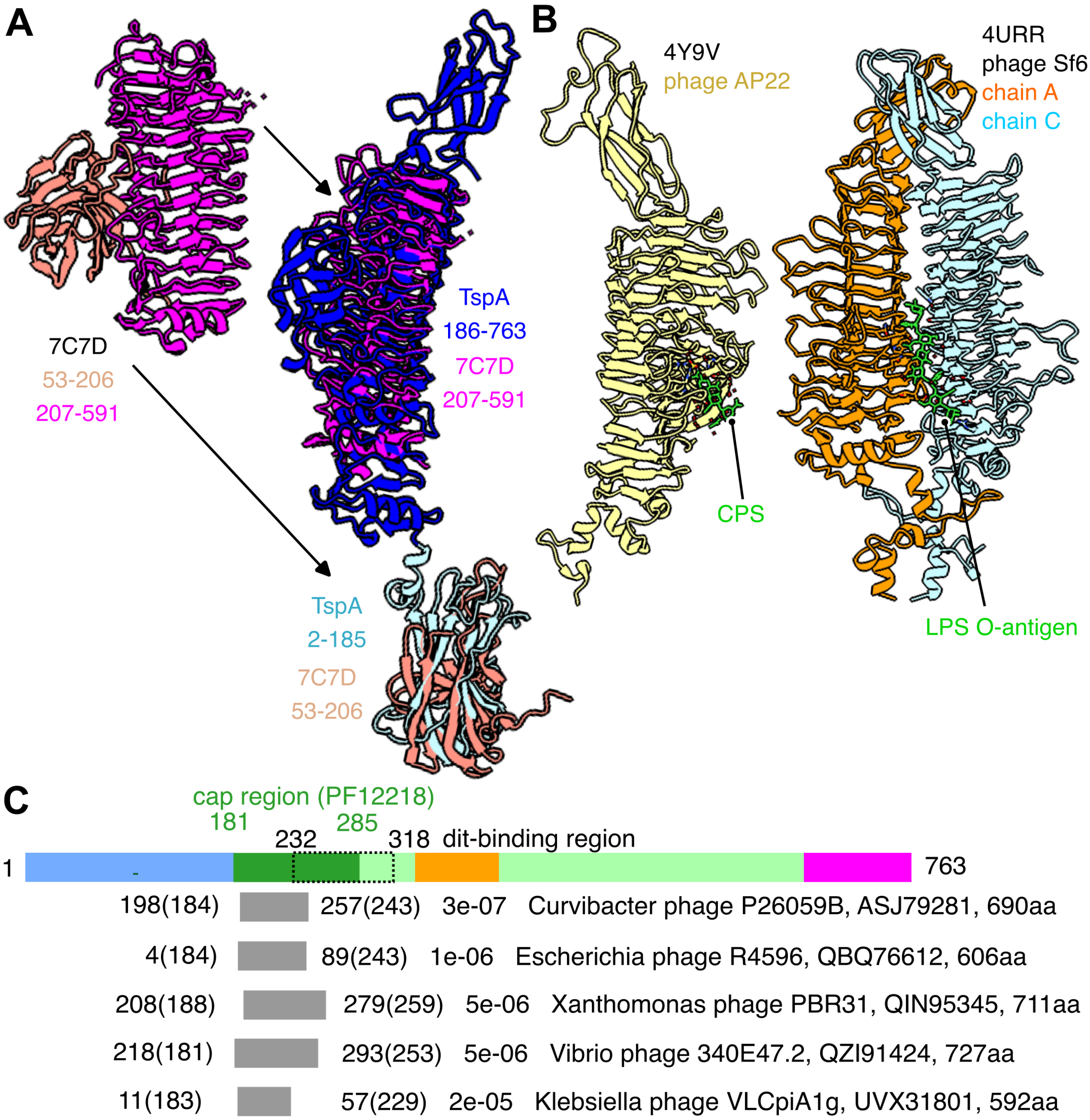


**Fig. S5. Comparison of TspA with other proteins in databases. A)** Structural alignment of *Streptomyces thermodiastaticus* α-1,3-glucanase (*2*) with TspA. On the left, a ribbon representation of the glucanase. The protein is coloured differently for the N-terminal β-sandwich (salmon) and central right-handed β-helix (magenta) domain. On the right, the superimposition of TspA on the glucanase is shown; the glucanase domains are superimposed separately. **B)** Ribbon representation of phage tailspike homologs of TspA whose structure was solved with a ligand (in green), binding to either intradomain (on the left) or inter-domain (on the right) cleft (*3*). The PDB codes of the structures are shown above. CPS, capsular polysaccharide; LPS, lipopolysaccharide. **C)** BLAST search with BLOSUM45 against the nr protein database of *Caudoviricetes* entries identifies hits only against the cap region. Top five hits are shown, with the residue range in the subject (and query in brackets), E-value, taxonomy name, subject protein code and length shown from left to right. The colour coding of the TspA is the same as in the main text, **Fig. 3**. A region binding the insertion domain of the distal tail (dit) protein is highlighted in a rectangle, with the delimiting residues depicted.

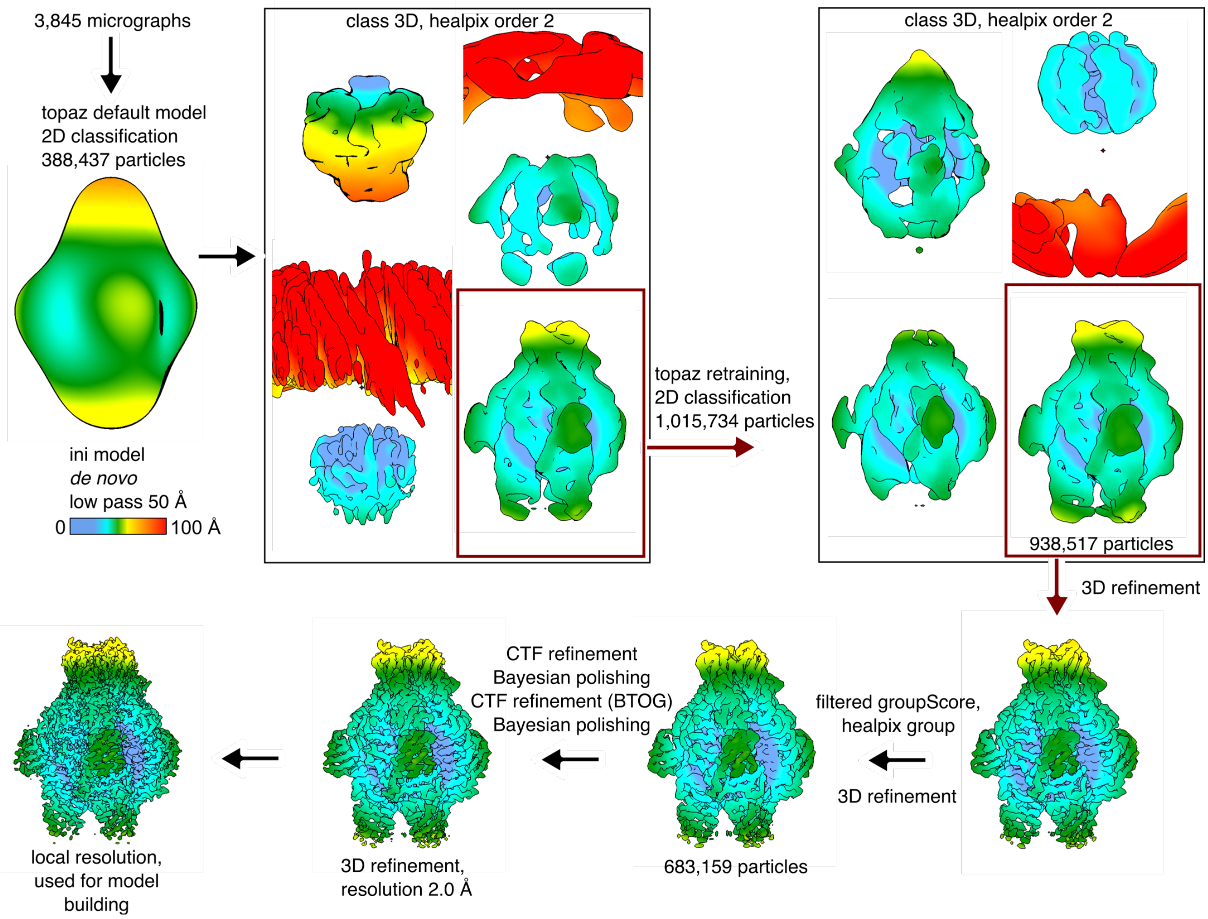


**Fig. S6. Single particle analysis of TspA incubated in buffer.** The analysis was done in RELION5 (*1*). The colour bar indicates the distance from the centre of the particle. The number of micrographs corresponds to the list filtered according to defocus, astigmatism and estimated resolution. The initial model was calculated *de novo*. Another 3D refinement was run after CTF refinement and Bayesian polishing. BTOG, beam tilt optics group. The star files were filtered and optics group edited using starpy scripts made by Dr Tibor Fuzik, CEITEC (https://github.com/fuzikt/starpy).

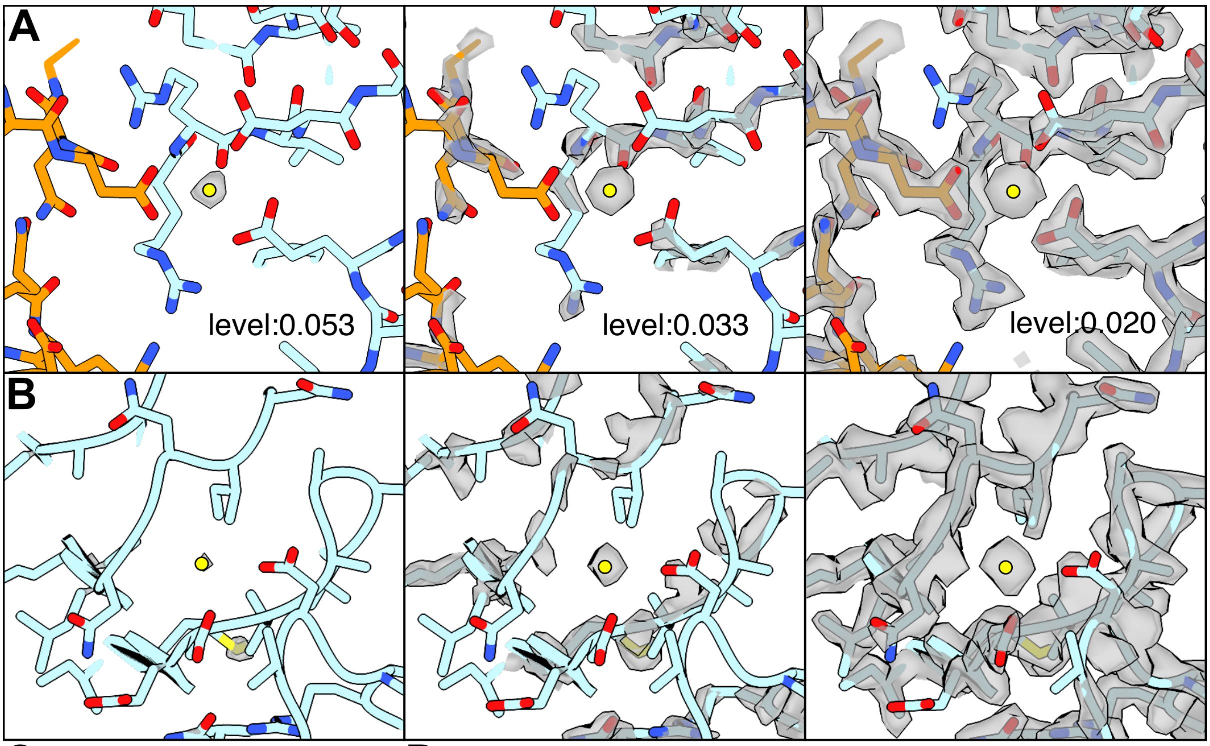


**C**

| Atoms | Scattering amplitudes [2 Å] |
|---|---|
| Na+ | 1.284 |
| Mg2+ | 1.472 |
| K+ | 2.575 |
| Ca2+ | 2.674 |
| S | 2.326 |
| C | 1.168 |
| N | 1.216 |
| O | 1.222 |

**D**

| Ca-Atom in TspA | Distance [Å] | Ca-Atom in PDB | Mean distance [Å]* |
|---|---|---|---|
| HOH/A | 2.43 | NA | NA |
| E271/Oε1 | 2.65 | Ca-Oε1 | 2.4 |
| E271/Oε2 | 2.42 | Ca-Oε2 | 2.6 |
| D287/Oδ1 | 2.58 | Ca-Oδ1 | 2.4 |
| D287/Oδ2 | 2.76 | Ca-Oδ2 | 2.6 |
| D299/Oδ2 | 2.54 | Ca-Oδ2 | 2.6 |
| R297/O | 2.4 | Ca-O (R) | 2.4 |
| HOH0/O | 2.79 | NA | NA |
| HOH1/O | 2.71 | NA | NA |
| HOH2/O | 2.56 | NA | NA |
| D542/Oδ1 | 2.74 | Ca-Oδ1 | 2.4 |
| D602/Oδ1 | 2.37 | Ca-Oδ1 | 2.4 |
| N635/O | 2.43 | Ca-O (R) | 2.4 |

**shown for the relevant distance peak*

**Fig. S7. Identification of calcium ions in the TspA map. A-B)** The map of TspA when incubated in YPS is shown in different density threshold levels. The ion (yellow circle) density shows at a higher threshold than the protein, implying its larger scattering amplitude. The inter-domain (A) and intra-domain (B) calcium coordination regions are shown. **C)** Scattering amplitudes of ions and atoms for the resolution of 2 Å, taken from International Tables for Crystallography (*4*). The ions with values higher than those of the C-O-N elements are highlighted in green. **D)** Ca-O distances observed in the structure, compared to the mean distances as estimated from <1.5 Å resolution structures in PDB (*5*).

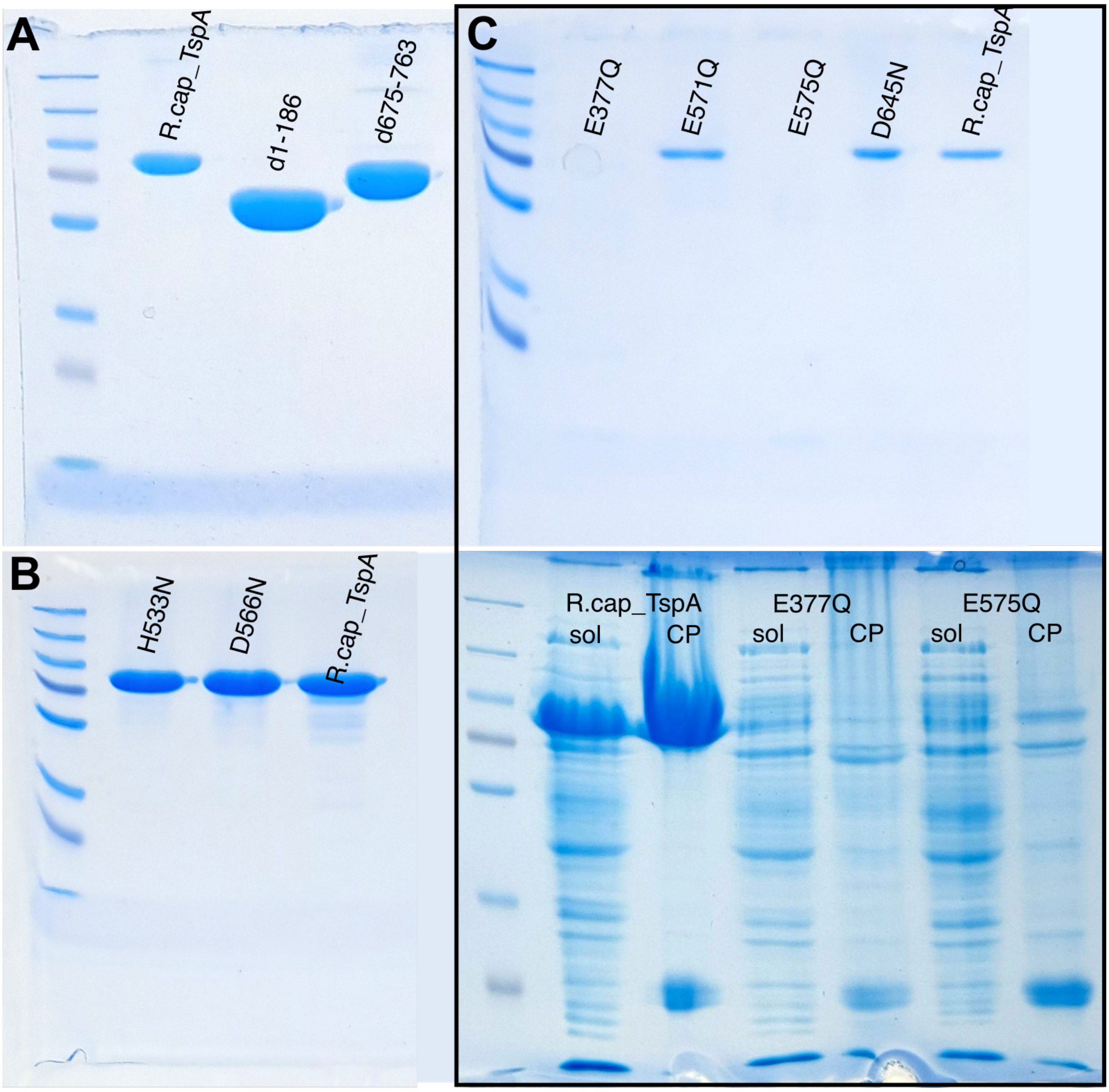


**Fig. S8. SDS-PAGE gels of purified mutant variants of TspA. A)** Truncated variants of TspA, compared to the wild-type protein. **B)** Mutants in the inter-domain active site. **C)** Mutants in the intra-domain active site. On the top, SEC-purified products are shown, with E377Q and E575Q showing no protein. On the bottom, crude purification of these variants, showing a weak band of the protein of expected size. The marker is PageRuler Plus Prestained Protein Ladder (10-250kDa, ThermoFisher Scientific).

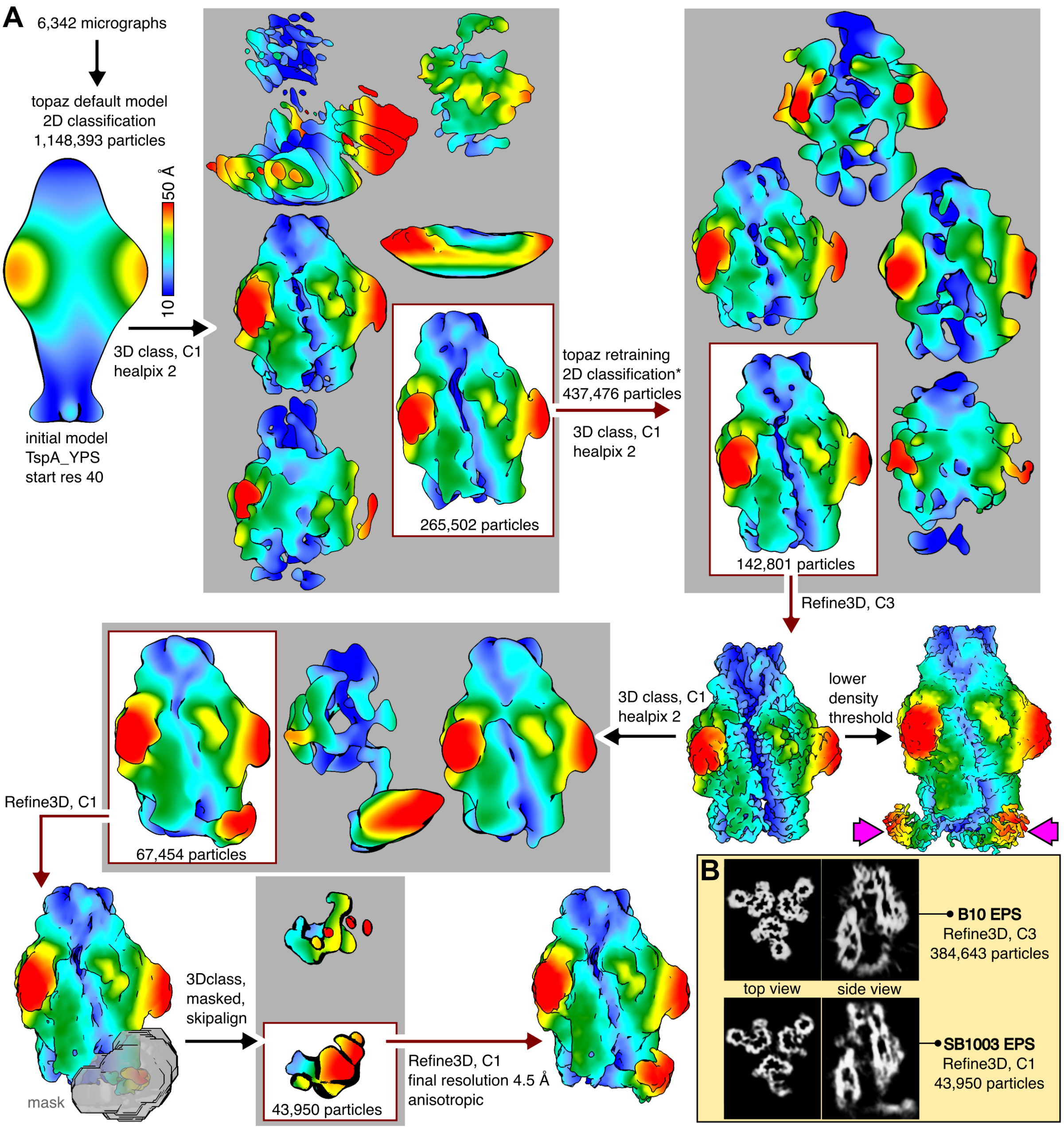


**Fig. S9. Single particle analysis of TspA incubated with a ligand. A)** TspA mixed with the resuspended biofilm of *R. capsulatus* SB1003. The analysis was done in RELION5 (*1*). The colour bar indicates the distance from the central z-axis of the particle. The initial model was taken from the dataset of TspA incubated in YPS. Magenta arrows point to extra density visible during initial C3 reconstruction, suggesting resolved N-terminal domains. *During this 2D classification, all particles were taken from class averages, which looked like side views; 1500 particles were taken from classes that looked like top views, leading to a less anisotropic reconstruction. **B)** Comparison of maps obtained from datasets on TspA mixed with extracellular polymeric substance (EPS) formed by *R. capsulatus* B10 and SB1003. Even in C3 symmetry, the dataset with B10 EPS showed more severe anisotropy.

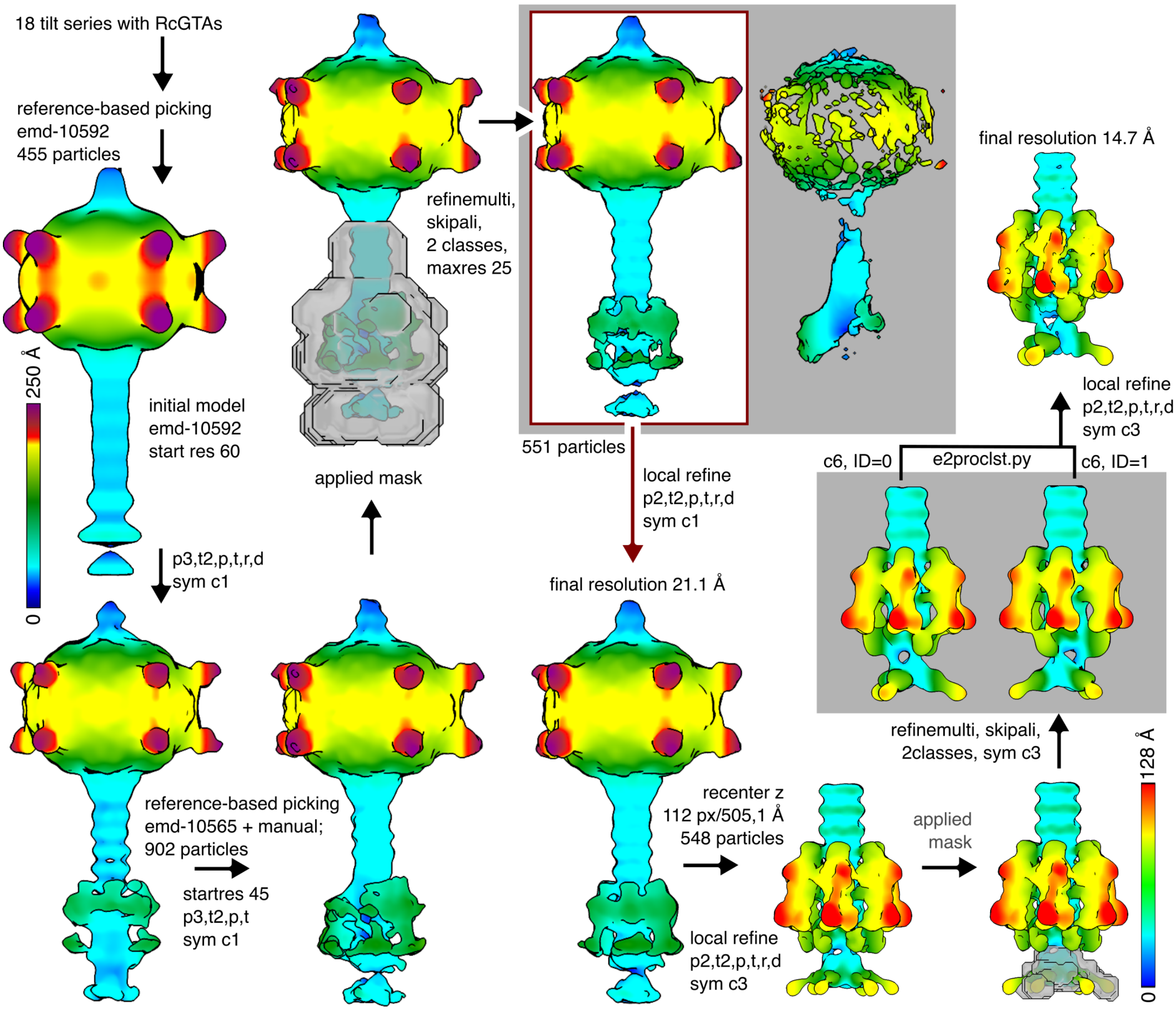


**Fig. S10. Subtomogram averaging pipeline of RcGTA particles imaged *in situ*.** The analysis was done in EMAN2 (*6*). The colour bar indicates the distance from the centre of the particle, the top one for the capsid, the bottom for the baseplate. The number of tilt series corresponds to those that aligned well and contained RcGTA particles.

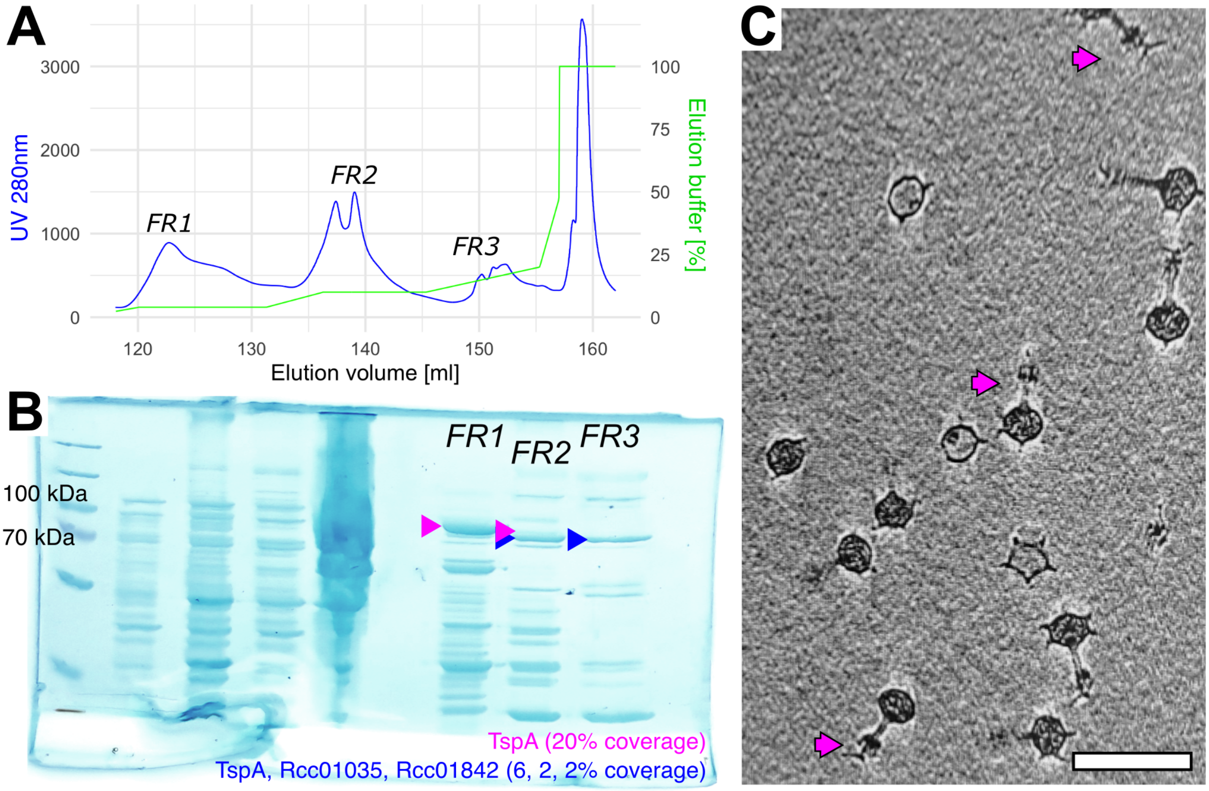


**Fig. S11. The alteration of the ion-exchange purification step to obtain RcGTAs with attached TspA.** A) Chromatogram of the ion-exchange chromatography, which was performed in a buffer containing calcium (20mM HEPES, pH 7.0; 5 mM CaCl2; NaCl equilibration-elution range from 0.01 to 2 M). The elution strategy was deduced from the results shown in **Fig. S2**, where fraction (FR)1 corresponded to the elution buffer concentration of the first diminished band of *Δ280ΔtspA*; FR2 to the second diminished band of *Δ280ΔtspA;* and FR3 to that expected for RcGTAs (*7*). B) SDS-PAGE of FR1-3. FR2 showed a similar band pattern to FR3, except for a band highlighted with a magenta arrow, which migrated similarly to the most abundant component of FR1 and corresponded to the expected size of TspA. MS analysis of selected bands (magenta, blue arrow) of FR2 confirmed its presence (protein identity shown at the bottom). Other lanes of the gels are not relevant for this study. C) Cryo-ET screening of the FR2 sample confirmed the presence of a bulky density at baseplates of RcGTAs (magenta arrows). Scale bar is 100 nm.

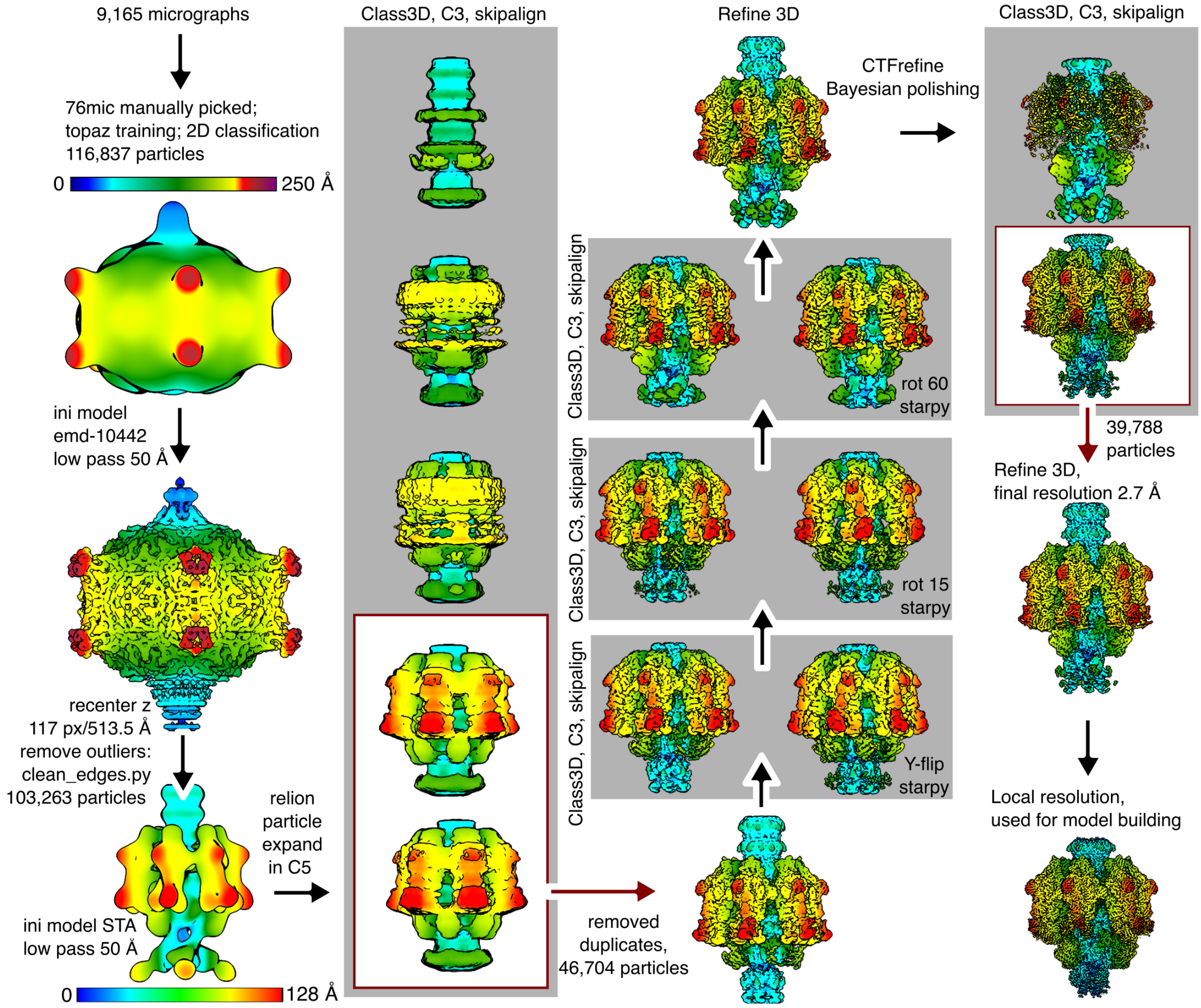


**Fig. S12. Single particle analysis of the RcGTA particle with attached TspA hexamer.** The analysis was done in RELION5 (*1*). The colour bar indicates the distance from the centre of the particle, the top one for the capsid, the bottom for the baseplate. The number of micrographs corresponds to the list filtered by defocus, astigmatism, and estimated resolution. The initial models used are shown in the low-pass resolution. Another 3D refinement was run after CTF refinement and Bayesian polishing. The star files were filtered to remove particles that would lie outside the micrograph after the recentering using the clean_edges.py script made by Dr Huw Jenkins, University of York (https://github.com/huwjenkins/em_scripts); and edited for rotating/Y-flipping of particles using starpy scripts made by Dr Tibor Fuzik, CEITEC (https://github.com/fuzikt/starpy). The two classes with disturbed TspA density from the initial classification were reprocessed and a class with well-resolved TspA density was identified; however, the addition of these particles to the final particle subset did not lead to a better-resolved map.

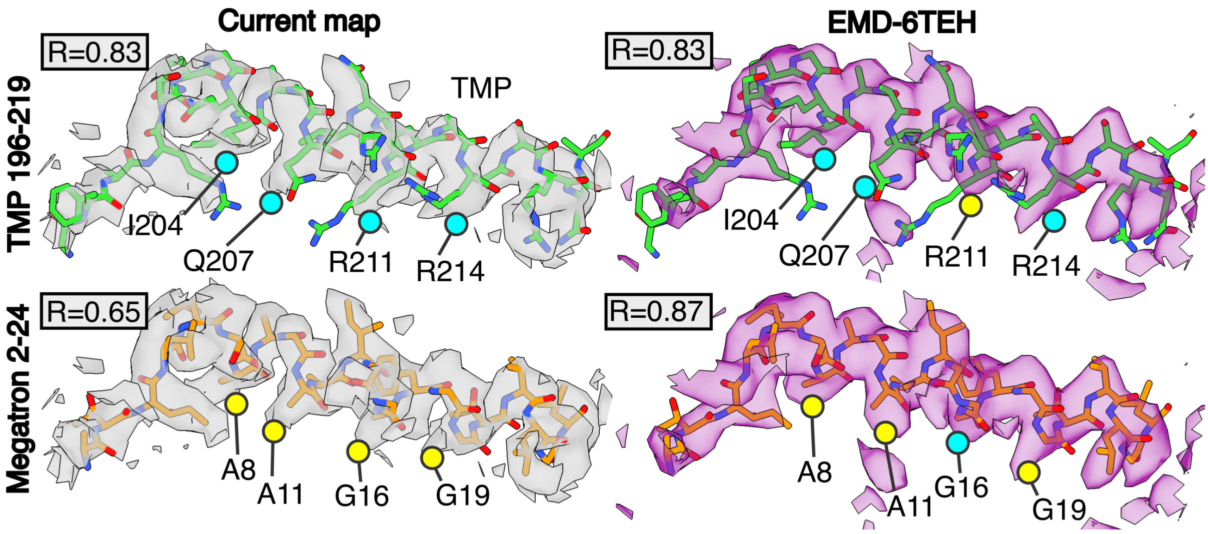


**Fig. S13. Reannotation of the central α-helix present in the cavity of the RcGTA baseplate.** The improved map obtained in this study (left) showed that the C-terminal part of the tape measure protein (TMP) fits the density better than the N-terminal part of megatron protein, which was previously proposed to form this helix (*7*). Four residues of the regions which differ in the size of the sidechain density are highlighted. In the current map (left), all four residues of TMP fitted better, while in the previously derived lower resolution map (right), three TMP residues fitted better; the density for R211 was not tractable. Cross-correlation coefficient (R) was calculated by comparing the experimental maps with maps simulated from the molecules in the same resolution using the fitmap command in ChimeraX (*8*).

**Table S1. LC-MS/MS analysis of a concentrated supernatant from the RcGTA-overproducing culture of *R. capsulatus* DE442.** Top 10 hits based on the normalised spectral counts are shown; structural proteins of RcGTA are shown in green.

| Accession | Score | Mass | No. Spectra | Norm spectral counts | No. Peptides | emPAI | Mol% | Description |
|---|---|---|---|---|---|---|---|---|
| A0A0E2PA72 | 6594 | 82144 | 315 | 4.70 | 42 | 6.82 | 0.84 | Tailspike TspA (Rcc02623) |
| A0A0E2PDC8 | 6122 | 42144 | 284 | 4.24 | 17 | 5.23 | 0.64 | Major capsid protein (Rcc01687) |
| A0A0E2PE20 | 4140 | 52495 | 206 | 3.07 | 38 | 16.89 | 2.08 | Glutamine synthetase |
| A0A0Q0R2P0 | 3111 | 82213 | 170 | 2.54 | 24 | 2.16 | 0.27 | Uncharacterized protein |
| A0A0E2PBE2 | 3547 | 57682 | 157 | 2.34 | 44 | 23.78 | 2.93 | 60 kDa chaperonin |
| A0A0E2PFC5 | 3794 | 138941 | 156 | 2.33 | 61 | 4.38 | 0.54 | Megatron (Rcc01698) |
| A0A0E2PJA9 | 3994 | 38170 | 146 | 2.18 | 19 | 4.78 | 0.59 | Tail fibre (Rcc00171) |
| A0A0E2PI29 | 2926 | 32489 | 99 | 1.48 | 6 | 1.79 | 0.22 | Porin |
| A0A0E2PCV2 | 2131 | 56258 | 92 | 1.37 | 30 | 9.32 | 1.15 | Uncharacterized protein |
| A0A0E2PDK7 | 1610 | 47139 | 83 | 1.24 | 20 | 5.37 | 0.66 | Portal (Rcc01684) |

**Table S2. Proteins with sequence and structural similarities to various domains of TspA.** The foldseek search (*9*) was done using an iterative mode. The table continues on the next page.

| HHpred (whole protein, top 10 hits) | | | | | | | | | | |
|---|---|---|---|---|---|---|---|---|---|---|
| pdb code | chain | residue range | Scientific Name | Function | E value | Position in query | Probability | Score | Ligand | Position in TspA |
| 8YK2 | B | 17-634 | *Bifidobacterium bifidum* JCM 1254 | Alpha-galactosidase | 1.20E-15 | 185-669 | 99.79 | 182.08 | no | 185-669 |
| 7JWF | C | 14-606 | *Pseudoalteromonas distincta* | Glycosid hydrolase family 110 | 1.60E-15 | 175-675 | 99.78 | 178.41 | alpha-(1,3)-galactobiose | 175-675 |
| 4RU5 | C | 3-578 | *Pseudomonas* phage phi297 | Tailspike gp27, | 6.50E-15 | 181-757 | 99.76 | 172.83 | no | 181-757 |
| 5ZRU | A | 232-564 | *Bacillus circulans* | Alpha-1,3-glucanase | 9.90E-15 | 154-672 | 99.75 | 170.39 | no | 154-672 |
| 3ZPP | A | 16-435 | *Streptococcus pneumoniae* | Cell wall surface anchor family protein | 1.30E-14 | 181-671 | 99.71 | 161.38 | no | 181-671 |
| 5LW3 | A | 1-360 | *Azotobacter vinelandii* | C-5 Epimerase | 4.40E-14 | 187-669 | 99.71 | 154.13 | no | 187-669 |
| 3GQ9 | A | 13-425 | *Bacillus* phage phi29 | Preneck appendage protein | 2.00E-13 | 179-670 | 99.69 | 160.57 | no | 179-670 |
| 2UVF | A | 152-607 | *Yersinia enterocolitica* | Exopolygalacturonase GH28 | 1.10E-14 | 184-667 | 99.68 | 171.26 | digalaturonic acid | 184-667 |
| 4MR0 | B | 108-576 | *Streptococcus pneumoniae* | Plasmin and fibronectin-binding protein A | 5.30E-13 | 181-668 | 99.65 | 158.28 | no | 181-668 |
| 5OLP | B | 38-452 | *Bacteroides thetaiotaomicron* DSM 2079 | Pectate lyase | 4.20E-13 | 182-670 | 99.63 | 151 | no | 182-670 |
| Foldseek (whole protein, top 5 hits) | | | | | | | | | | |
| pdb code | chain | residue range | Scientific Name | Function | E value | Position in query | Probability | Score | Ligand | Position in TspA |
| 2VJI | A | 24-598 (598) | *Escherichia* phage HK620 | tailspike | 2.43E-49 | 159-760 (762) | 1 | 954 | no | 160-761 (763) |
| 5LW3 | A | 2-373 (381) | *Azotobacter vinelandii* | Mannuronan C-5 epimerase module | 1.71E-23 | 188-674 (762) | 1 | 402 | no | 189-675 (763) |
| 4mr0 | A | 2-444 (456) | *Streptococcus pneumoniae* | surface adhesin PfbA | 2.81E-09 | 187-696 (762) | 1 | 237 | no | 188-670 (763) |
| 3GQ7 | A | 19-454 (605) | *Bacillus* phage phi29 | tailspike | 1.25E-10 | 188-667 (762) | 1 | 231 | no | 189-668 (763) |
| 7C7D | B | 4-524 (539) | *Streptomyces thermodiastaticus* | Alpha-1,3-glucanase | 3.69E-11 | 37-669 (762) | 1 | 220 | no | 38-670 (763) |
| Foldseek (N-terminal domain, top 3 nonvirus hits plus top 5 virus hits) | | | | | | | | | | |
| pdb code | chain | residue range | Scientific Name | Function | E value | Position in query | Probability | Score | Ligand | Position in TspA |
| 4XUO | B | 1-153 (156) | *Paenibacillus bacrinonensis* | xylan-binding domain CBM22-1 | 8.31E-06 | 8-176 (184) | 1 | 183 | no | 9-177 (763) |
| 2ZEW | B | 5-147 (147) | *Caldanaerobius polysaccharolyticus* | cabohydrate binding module Family 16 | 2.19E-05 | 8-176 (184) | 1 | 172 | no | 9-177 (763) |
| 2WZE | B | 5-146 (516) | *Acetivibrio thermocellus* | cabohydrate binding module CBM22-1 | 7.96E-05 | 10-173 (184) | 1 | 172 | tribeta-D-xylopyranose-(1-4) | 11-174 (763) |
| 5W6H | C | 551-697 (697) | *Kuttervirus* CBA120 | tailspike C-terminal domain | 6.70E-05 | 8-176 (184) | 1 | 167 | no | 9-177 (763) |
| 7XYC | A | 421-562 (562) | *Klebsiella* phage Kp7 | tailspike C-terminal domain | 5.18E-05 | 8-176 (184) | 1 | 158 | no | 9-177 (763) |
| 2GSY | P | 195-327 (430) | Infectious bursal disease virus | capsid protrusion | 2.03E-02 | 40-176 (184) | 0.97 | 76 | no | 41-177 (763) |
| 5E7T | B | 52-183 (286) | *Lactococcus* phage Tuc2009 | minor structural protein 5 | 2.81E-02 | 51-174 (184) | 0.93 | 70 | no | 52-175 (763) |
| 6QCC | B | 67-171 (182) | Broad bean stain virius | capsid protrusion | 6.56E-03 | 76-176 (184) | 0.87 | 65 | no | 77-177 (763) |

| Foldseek (C-terminal domain, top 3 nonvirus hits plus virus hits) | | | | | | | | | | |
|---|---|---|---|---|---|---|---|---|---|---|
| pdb code | chain | residue range | Scientific Name | Function | E value | Position in query | Probability | Score | Ligand | Position in TspA |
| 7Z28 | A | 35-105 (859) | *Homo Sapiens* ERAP1 | M1 zinc aminopeptidase | 7.98E-01 | 4-83 (89) | 0.25 | 42 | di-2-acetamido-2-deoxy-beta-D-glucopyranos | 678-757 (763) |
| 4A3Z | A | 41-119 (136) | *Clostridium perfringens* | Alpha-N-Acetylglucosaminidase | 8.46E-01 | 2-89 (89) | 0.16 | 37 | no | 676-763 (763) |
| 8EO2 | B | 182-258 (271) | *Lutzomyia longipalpis* | Lufaxin | 2.45E+00 | 3-83 (89) | 0.16 | 37 | di-2-acetamido-2-deoxy-beta-D-glucopyranos | 677-757 (763) |
| 6CL6 | B | 293-361 (364) | *Pseudomonas aeruginosa* R2 pyocin | C-terminal domain of a fibre | 1.53E+00 | 4-83 (89) | 0.44 | 49 | no | 678-757 (763) |
| 7EEA | C | 579-644 (646) | Cyanophage Pam1 | TSP insertion domain | 2.60E+00 | 3-83 (89) | 0.15 | 36 | no | 677-757 (763) |
| 8ECI | 1 | 9-77 (124) | *Anthrobacter* phage Bridgette | Decoration protein | 2.93E+00 | 3-87 (89) | 0.07 | 27 | no | 677-761 (763) |
| 1UF2 | D | 189-289 (417) | Rice dwarf virus | Outer capsid protein P8 | 6.69E+00 | 4-83 (89) | 0.04 | 23 | no | 678-757 (763) |

*tailspikes are shown in green*

**Table S3. The analysis of interfaces in the RcGTA baseplate structure.** The values were calculated using PDBePISA v1.52 (*10*).

| Interface | Chains | Burried surface area [Å²] | Energy [kcal/mol] |
|---|---|---|---|
| | D vs B | 425.7 | -2.2 |
| | A vs E' | 372.5 | -2.4 |
| | C vs D | 104.7 | -0.4 |
| TspA trimer:TspA trimer | A vs F' | 98.2 | -0.1 |
| | B vs E' | 52.8 | -0.1 |
| | E vs B | 47.3 | -0.3 |
| | **Sum** | **1101.2** | **-5.5** |
| TspA trimer:Distal tail | B vs I | 574.2 | 0.7 |
| | A vs L | 570.6 | -2.1 |
| | B vs L | 119.3 | 1.3 |
| | B vs K | 35.7 | 0.2 |
| | **Sum** | **1299.8** | **0.1** |
| TspA trimer:Tail tube | B vs G | 209.2 | -0.8 |
| | B vs H | 29.4 | -0.3 |
| | B vs J | 10.7 | 0 |
| | **Sum** | **249.3** | **-1.1** |
| TspA trimer:Megatron | C vs N | 156.3 | -0.5 |
| | **Sum** | **156.3** | **-0.5** |
| Distal tail:Tail tube | K vs J | 503.3 | -8.9 |
| | K vs I' | 103.9 | -2.4 |
| | K vs I | 12.3 | -0.4 |
| | **Sum** | **619.5** | **-11.7** |
| Tail tube:Tail tube | I vs H | 432.7 | -3.1 |
| | I vs G | 147 | -1 |
| | I vs G' | 82.8 | -1.5 |
| | **Sum** | **662.5** | **-5.6** |
| Megatron:Distal tail | N vs K' | 307.8 | -5.1 |
| | N vs L | 60.4 | -0.8 |
| | **Sum** | **368.2** | **-5.9** |
| Hub:Distal tail | M vs L | 413.9 | -5.2 |
| | M vs K | 416.4 | 1.4 |
| | M vs L' | 91.4 | -1.8 |
| | **Sum** | **921.7** | **-5.6** |
| | D2 vs J4 | 413.2 | -2.5 |
| | D2 vs K4 | 126.8 | -1.1 |
| Spike base:capsid (6TB9) | D2 vs I4 | 116.2 | -1.2 |
| | D2 vs L4 | 12.9 | 0.3 |
| | **Sum** | **669.1** | **-4.5** |
| | B vs F | 876.2 | -8.9 |
| | B vs E | 676.1 | -1.7 |
| | B vs D | 154.3 | -1.6 |
| Tail fibre:Megatron | A vs E | 112.2 | -1.6 |
| | A vs D | 65.2 | 0.1 |
| | **Sum** | **1884** | **-13.7** |

*the symmetry expanded version of a chain is depicetd by apostrophe*

**Table S4. Bioinformatic analysis of the RcGTA distal tail protein**. TspA-encoding GTA species as shown in **Fig. 1D**, and top phage hits are shown, as identified using blastp (*11*). The identity columns are coloured in descending fashion using a palette green-yellow-orange.

| Species | Whole protein | | | ID (89-174) | |
|---|---|---|---|---|---|
| | Query [%] | Identity [%] | E-value | Query [%] | Identity [%] |
| *Paracoccus* GTA | 100 | 72.73 | 9E-112 | 97 | 54.76 |
| *Ruegeria* GTA | 100 | 69.23 | 1E-102 | 100 | 52.87 |
| *Phaeobacter* GTA | 100 | 65.87 | 2E-102 | 100 | 48.28 |
| *Dinoroseobacter* GTA | 100 | 70.19 | 4E-102 | 99 | 55.81 |
| *Rhodovulum* GTA | 100 | 69.23 | 6E-102 | 99 | 53.49 |
| *Sulfitobacter* GTA | 100 | 65.38 | 7E-102 | 100 | 52.87 |
| *Ketogulonicigenium* GTA | 100 | 62.02 | 6E-96 | 99 | 44.19 |
| *Roresobacter* phage RDJL3 | 100 | 54.81 | 3E-79 | 99 | 41.86 |
| *Rhodobacter* phage RcRudolph | 100 | 54.81 | 3E-76 | 94 | 48.78 |
| *Rhodobacter* phage RcKemmy | 100 | 53.37 | 3E-75 | 99 | 51.16 |
| *Rhodobacter* phage RcSimone-Hastad | 100 | 53.85 | 2E-71 | 91 | 48.10 |
| *Dinoroseobacter* phage R26L | 100 | 50.24 | 3E-70 | 99 | 44.19 |
| *Paracoccus* phage R3 | 100 | 48.80 | 2E-69 | 91 | 43.04 |
| *Rhodobacter* phage RcOceanus | 100 | 50.48 | 8E-69 | 94 | 46.34 |
| *Rhodobacter* phage RcFrancesLouise | 100 | 50.48 | 9E-69 | 94 | 46.34 |
| *Rhodobacter* phage RcBaka | 100 | 50.48 | 3E-68 | 91 | 46.84 |
| *Desulfofustis* phage LS06-2028-MD02 | 100 | 51.20 | 2E-65 | 77 | 40.30 |
| *Rhodobacter* phage RcCronus | 100 | 52.07 | 4E-65 | 91 | 48.10 |
| *Rhodobacter* phage RcRhea | 100 | 51.61 | 5E-65 | 91 | 48.10 |
| *Rhodobacter* phage RcZahn | 100 | 45.19 | 2E-61 | 91 | 43.04 |
| *Dinoroseobacter* phage R5C | 89 | 49.73 | 1E-59 | 95 | 45.78 |

*ID, insertion domain*

**Table S5. Primers used in this study.**

| Primer name | Sequence (5'-3') | Amplified gene/Mutation |
|---|---|---|
| RcTspA_F | TCCAGGGACCAGCAATGAACATCGCGATCACCGATG | *R. capsulatus* TspA |
| RcTspA_R | TGAGGAGAAGGCGCGCCCGCGGATCTGACCTGAG | |
| RmTspA_F | TCCAGGGACCAGCAATGAACAAGGCAATCACCG | *R. mobiis* TspA |
| RmTspA_R | TGAGGAGAAGGCGCGCTAGCGGTCGATGCGGA | |
| PpTspA_F | TCCAGGGACCAGCAATGAACATAGCGATCACTGG | *P. piscinae* TspA |
| PpTspA_R | TGAGGAGAAGGCGCGTTACTTGTCCACCCGGAC | |
| E337Q_F | GGCCGCCAGGTCTATGTGCTGGCGGTCGATC | TspA_E337Q |
| E337Q_R | ATAGACCTGGCGGCCGACGCCGGTGCCGCTC | |
| H533Q_F | CAACCAATGGTTCCAGGGCGACGGGCAG | TspA_H533Q |
| H533Q_R | TGGAACCATTGGTTGCCCACGATCACATGG | |
| D566N_F | TACATCAACAACAACTTCATCGAATGGACG | TspA_D566N |
| D566N_R | GTTGTTGTTGATGTAATTGCCGGTCACC | |
| E571Q_F | TTCATCCAATGGACGAACGAATATTC | TspA_E571Q |
| E571Q_R | CGTCCATTGGATGAAGTTGTTGTCGATG | |
| E575Q_F | ACGAACCAATATTCGCCCGATCCGAACCTCG | TspA_E575Q |
| E575Q_R | CGAATATTGGTTCGTCCATTCGATGAAGTTG | |
| D645N_F | AAGGTGAACACGAGCTTTGCCGATCTCGAC | TspA_D645N |
| D645N_R | GCTCGTGTTCACCTTCTCGATCCGGTTGATG | |
| M186_F | TCCAGGGACCAGCAATGGATGTGGTCGATGTGC | TspA_d1-185 |
| M186_R | TGAGGAGAAGGCGCGCCCGCGGATCTGACCTGAG | |
| D675_F | TCCAGGGACCAGCAATGAACATCGCGATCACCGATG | TspA_d675-763 |
| D675_R | TGAGGAGAAGGCGCGTCAGTTCACCGTCATCTGGCTC | |

### Table S6: Cryo-EM and ET data collection and processing information.

| Sample | Single particle analysis | | | | Subtomogram averaging |
|---|---|---|---|---|---|
| | TspA in YPS | TspA in buffer | TspA with ligand | RcGTA baseplate | RcGTAs produced *in situ* |
| **Data collection parameters** | | | | | |
| Magnification | ×165,000 | ×165,000 | ×165,000 | ×81,000 | ×19,500 |
| Pixel size [Å] | 0.743 | 0.743 | 0.743 | 1.10 | 4.55 |
| Camera (energy filter) [manufacturer] | Falcon 4 (Selectris X) [TFS] | Falcon 4 (Selectris X) [TFS] | Falcon 4 (Selectris X) [TFS] | K3 (BioQuantum) [Gatan] | K3 (BioQuantum) [Gatan] |
| Slit size [kV] | 10 | 10 | 10 | 20 | 20 |
| Instrument* | Krios Alpha, SLAC-Stanford | Krios Alpha, SLAC-Stanford | Krios Alpha, SLAC-Stanford | Krios Beta, SLAC-Stanford | Krios Beta, SLAC-Stanford |
| Voltage [kV] | 300 | 300 | 300 | 300 | 300 |
| Data collection software | EPU | EPU | EPU | EPU | Tomo5 |
| Total exposure dose [$e^-/Å^2$] | 39.31 | 39.7 | 46.96 | 28.542 | 119.88 |
| Exposure dose per tilt [$e^-/Å^2$] | 39.31 | 39.7 | 46.96 | 28.542 | 3.24 |
| Raw image format (number of frames) | tiff (45) | tiff (37) | tiff (50) | tiff (30) | mrcs (4) |
| Data collection scheme | single acquisition | single acquisition | single acquisition | single acquisition | Bi-directional, starting tilt 0° |
| Number of tilts | 1 | 1 | 1 | 1 | 37 |
| Tilt span [°] | NA | NA | NA | NA | ±54 |
| Tilt increment [°] | NA | NA | NA | NA | 3 |
| Number of acquired micrographs/tilt series | 4,604 | 4,355 | 6,342 | 10,951 | 83 |
| Number of used micrographs/tilt series | 3,605 | 3,845 | 5,731 | 9,165 | 18 |
| Targeted defocus [µm] | 0 - 1.6 | 0 - 1.6 | 0.2-1.6 | 0.2 - 2 | 4 |
| **Data processing parameters** | | | | | |
| Reconstruciton software | RELION5 | RELION5 | RELION5 | RELION5 | EMAN2 |
| Symmetry | C3 | C3 | C1 | C3 | c1 whole capsid. c3 baseplate |
| Particle picking method | topaz neural network | topaz neural network | topaz neural network | topaz neural network | reference-based, EMD-10565 |
| Initial number of particles† | 354,994 | 1,015,734 | 437,468 | 116,837 | 902 |
| Final number of particles | 82,697 | 683,159 | 43,950 | 39,788 | 551 |
| Initial model | from TspA in buffer | *de novo* | from TspA in YPS | EMD-10442; STA base | EMD-10592 |
| Map resolution [Å] | 2.0 | 2.0 | 4.5‡ | 2.7 | c1=21.1, c3=14.7 |
| FSC threshold | 0.143 | 0.143 | 0.143 | 0.143 | 0.2 |

**all manufactured by TFS; † estimated after first 2D classification; ‡ anisotropic map; TFS, Thermo Fisher Scientific*

**Supplementary references:**


1. D. Kimanius, L. Dong, G. Sharov, T. Nakane, S. H. W. Scheres, New tools for automated cryo-EM single-particle analysis in RELION-4.0. *Biochem J* **478**, 4169–4185 (2021).

2. T. Itoh, N. Panti, J. Hayashi, Y. Toyotake, D. Matsui, S. Yano, M. Wakayama, T. Hibi, Crystal structure of the catalytic unit of thermostable GH87 α-1,3-glucanase from *Streptomyces thermodiastaticus* strain HF3-3. *Biochemical and Biophysical Research Communications* **533**, 1170–1176 (2020).

3. Y. Kang, U. Gohlke, O. Engström, C. Hamark, T. Scheidt, S. Kunstmann, U. Heinemann, G. Widmalm, M. Santer, S. Barbirz, Bacteriophage Tailspikes and Bacterial O-Antigens as a Model System to Study Weak-Affinity Protein–Polysaccharide Interactions. *J. Am. Chem. Soc.* **138**, 9109–9118 (2016).

4. C. Colliex, J. M. Cowley, S. L. Dudarev, M. Fink, J. Gjønnes, R. Hilderbrandt, A. Howie, D. F. Lynch, L. M. Peng, G. Ren, A. W. Ross, V. H. Smith, J. C. H. Spence, J. W. Steeds, J. Wang, M. J. Whelan, B. B. Zvyagin, Electron diffraction. 259–429 (2006).

5. M. Bazayeva, C. Andreini, A. Rosato, A database overview of metal-coordination distances in metalloproteins. *Acta Crystallogr D Struct Biol* **80**, 362–376 (2024).

6. M. Chen, J. M. Bell, X. Shi, S. Y. Sun, Z. Wang, S. J. Ludtke, A complete data processing workflow for cryo-ET and subtomogram averaging. *Nat Methods* **16**, 1161–1168 (2019).

7. P. Bárdy, T. Füzik, D. Hrebík, R. Pantůček, J. Thomas Beatty, P. Plevka, Structure and mechanism of DNA delivery of a gene transfer agent. *Nat Commun* **11**, 3034 (2020).

8. E. F. Pettersen, T. D. Goddard, C. C. Huang, E. C. Meng, G. S. Couch, T. I. Croll, J. H. Morris, T. E. Ferrin, UCSF ChimeraX: Structure visualization for researchers, educators, and developers. *Protein Sci* **30**, 70–82 (2021).

9. M. van Kempen, S. S. Kim, C. Tumescheit, M. Mirdita, J. Lee, C. L. M. Gilchrist, J. Söding, M. Steinegger, Fast and accurate protein structure search with Foldseek. *Nat Biotechnol* **42**, 243–246 (2024).

10. E. Krissinel, K. Henrick, Inference of Macromolecular Assemblies from Crystalline State. *Journal of Molecular Biology* **372**, 774–797 (2007).

11. S. F. Altschul, T. L. Madden, A. A. Schäffer, J. Zhang, Z. Zhang, W. Miller, D. J. Lipman, Gapped BLAST and PSI-BLAST: a new generation of protein database search programs. *Nucleic Acids Research* **25**, 3389–3402 (1997).
